# Redox Regulation of Cell Migration via Nischarin S-glutathionylation

**DOI:** 10.1101/2025.09.25.678565

**Authors:** Madhu C. Shivamadhu, Dhanushika S. K. Kukulage, Rayavarapu Padmavathi, Daniel Oppong, Faezeh Mashhadi Ramezani, Denaye N. Eldershaw, Brett M. Collins, Young-Hoon Ahn

**Author notes:** Corresponding author: Young-Hoon Ahn.

## Abstract

Reactive oxygen species (ROS) are central players in redox signaling, controlling all biological processes in human health. Many reports demonstrated that ROS play essential roles in regulating cell migration and invasion, while contributing to cancer progression and metastasis, potentially via inducing protein cysteine oxidations. Nevertheless, specific redox players involved in cell migration and invasion remain ill-defined. In this report, we found that Nischarin (NISCH), established as a tumor suppressor, is susceptible to S-glutathionylation, selectively at Cys185 located near its leucine-rich repeat (LRR) domains, which are implicated in protein-protein interactions with Rac1 and PAK1. We demonstrated that epithelial breast cancer cell lines, MCF7 and MDA-MB-231, expressing NISCH wild-type (WT), compared to its cysteine mutant (C185S), exhibit increased migration and invasion in response to oxidative stress, such as limited glucose. Mechanistically, NISCH S-glutathionylation reduced its binding to Rac1 and PAK1, without altering its binding to integrin α5. The dissociation of NISCH led to the activation of Rac1 and PAK1, resulting in the localization of Rac1 to the cell periphery, which facilitates lamellipodia formation. The activated PAK1 increased the phosphorylation of the LIMK1-cofilin axis, thereby further enhancing actin filament dynamics that promote cell migration. Based on the mechanistic analysis, we produced an engineered NISCH construct, composed of the N-terminal PX and LRR domains. We demonstrated that the engineered NISCH PX-LRR constructs, particularly one lacking the S-glutathionylation site (i.e., C185S), can suppress the migration, invasion, and colony formation of MDA-MB-231 cells, regardless of the presence of oxidative stress. Our data reports a new redox player in cell migration and invasion, while supporting the potential application of NISCH-derived protein-based therapeutics for breast cancer.

## Introduction

Cell migration and invasion play a central role in diverse physiological processes, including embryonic development, immune response, angiogenesis, and wound healing.^1^ In addition, dysregulation of cell migration and invasion is a prominent feature in pathologies, including cancer metastasis,^2^ inflammatory diseases (e.g., arthritis),^3^ and vascular diseases (e.g., atherosclerosis).^4^ Importantly, a large body of evidence indicates that cell migration and invasion are regulated by reactive oxygen species (ROS), hydrogen peroxide (H_2_O_2_), or redox signaling.^5,6^ For example, H_2_O_2_ acts as a chemokine directing the migration of leukocytes.^7^ H_2_O_2_ is locally produced at the polarized membrane of the migrating cell, mediating the formation of protrusions.^8^ Growth factors or integrin activation produce H_2_O_2_, leading to the formation of focal adhesions (via activating Src and FAK).^6,9^ H_2_O_2_ or oxidative stress activates Rho-GTPase signaling (e.g., Rac1, Cdc42, and RhoA),^10^ which generates traction force via cytoskeleton reorganization. H_2_O_2_ or oxidative stress promotes cell invasion by weakening cell-cell junctions and increasing the extracellular matrix degradation.^6^ Accordingly, high expressions of oxidases, including NADPH oxidases and lysyl oxidase, are reported in cancer cells, contributing to cancer migration, invasion, and metastasis.^11,12^

ROS play a role, in part, by inducing protein cysteine oxidation. Protein S-glutathionylation, a disulfide bond of protein cysteine with glutathione, is one of the major cysteine oxidations that occur in response to ROS and diverse chemical, biological, and medical factors,^13^ including electrophiles,^14^ metals,^15,16^ growth factors and cytokines,^17–19^ pathogens,^20^ nutrients,^21,22^ and radiation.^23^ Evidence continues to discover significant roles of S-glutathionylation in regulating physiology and diseases.^13,24^ Consistently, examples of S-glutathionylation in regulating cell migration and invasion have been reported. For example, previous studies have identified target proteins of S-glutathionylation regulating cell migration, including MAPK phosphatase-1 (MKP-1),^22^ low molecular weight protein tyrosine phosphatase (LMW-PTP),^25^ 14-3-3 zeta,^26^ and actin.^19^ In addition, we have recently demonstrated that glutathionylation of p120-catenin,^27^ the Ser/Thr phosphatase PP2Cα,^17^ and the lipid-binding protein FABP5^28^ regulates the migration and invasion of epithelial cancer cells. These examples underscore the importance of S-glutathionylation in regulating cell migration, adhesion, and invasion.

Over the years, multiple proteomic studies have profiled a large number of candidate cysteines susceptible to S-glutathionylation (n > 2,000),^13^ often the first step in discovering new redox-regulatory cysteines. Despite the large number of identified proteins, it remains challenging to uncover regulatory cysteines involved in specific biological processes, such as cell migration and invasion. To address this challenge, we previously developed an integrative strategy that combines proteomic databases, bioinformatics, and biological screening (i.e., cell migration),^17^ which led to the identification of three new proteins (i.e., PP2Cα, Nischarin, and ARHGEF7) that potentially regulate cell migration via S-glutathionylation.^17^ Among the three proteins, we reported the regulatory role of PP2Cα S-glutathionylation in epidermal growth factor (EGF)-induced cell migration.^17^

In this report, we demonstrate Nischarin (NISCH) S-glutathionylation and its functional effects in cell migration and invasion. NISCH is a large accessory and adaptor protein (human, amino acids 1-1504) that is ubiquitously expressed in all major tissues, interacting with protein partners in signaling pathways.^29^ NISCH is primarily established as a tumor suppressor, negatively affecting signaling proteins implicated in cancer cell migration, invasion, and metastasis.^29,30^ For example, NISCH was identified as a protein partner of integrin α5, negatively inhibiting integrin-dependent cell migration.^31^ In addition, NISCH binds and inhibits activities of PAK1 and Rac1,^32–34^ two pleiotropic oncogenes that drive cytoskeletal remodeling (e.g., lamellipodia), MAPK activation, and cell cycle progression.^35,36^ NISCH also directly binds to and inhibits LIMK1, an oncogenic effector of PAK1, thereby further inhibiting actin cytoskeletal dynamics.^37^ Consequently, NISCH expression reduces focal adhesion, motility, and epithelial-mesenchymal transition (EMT) in breast cancer cells.^38–40^ Consistently, the reduced level of NISCH expression correlates with aggressive stages of breast cancer.^29,41^ Beyond cancers, NISCH is highly expressed in the brain, including cortical neurons,^29^ where it suppresses neurite outgrowth and dendrite formation, potentially via inhibiting PAK1.^42^ Also, higher NISCH expression is associated with psychiatric disorders, such as Schizophrenia, autism, and depression.^43–45^ As well as PAK and Rac1, NISCH has been shown to be a late endosomal effector of RAB GTPases, including RAB9 and RAB14.^46^ Lastly, NISCH binds to insulin receptor substrate 4 (IRS4), increasing insulin signaling and regulating energy homeostasis.^29^ Despite these analyses, the redox regulation of NISCH has not been reported.

We demonstrate that NISCH is susceptible to S-glutathionylation selectively at Cys185 out of its 34 available cysteines. NISCH C185 S-glutathionylation disrupts its interaction with PAK1 and Rac1, thereby activating the LIMK1-cofilin downstream pathways and increasing cancer cell migration and invasion in oxidative stress. Building upon the mechanistic understanding, we developed and demonstrated that an engineered soluble NISCH construct (enNISCH C185S), which is devoid of the S-glutathionylation site, is effective in suppressing the migration and invasion of metastatic breast cancer cells, suggesting its potential use in therapeutic applications.

## Result

### NISCH is glutathionylated selectively at Cys185 in oxidative stress

To identify cysteine glutathionylation regulating cell motility, we previously searched for migration-associated proteins (n = 37) from a list of S-glutathionylated proteins (n = 1,351).^17^ Out of 37 proteins, we selected and analyzed 9 individual proteins (and 14 cysteines) in the in vitro migration assay. Among the 9 proteins, NISCH wild-type (WT) or its Cys mutant (C185S) caused differential migration levels of MDA-MB-231 cells in response to oxidative stress,^17^ suggesting that NISCH C185 oxidation potentially regulates cell migration.

NISCH comprises several distinct domains or regions at the N-terminus (amino acids 1-824), including phox homology (PX), leucine-rich repeat (LRR), coiled-coil (CC), and integrin-binding (ITGBD) domains, while its C-terminal regions (amino acids 825-1501) have less defined domains (Figure 1A).^47^ However, AlphaFold^48^ predicts that the C-terminal regions have three folded structures, and the last two regions are annotated as pleckstrin homology (PH) or PH-like domains (Pfam and UniProt)^47^ (Figure 1A). It is also notable that there are long intervening regions between domains (amino acids 428-586 and 1014-1106) that are predicted to be disordered and flexible. Interestingly, NISCH contains a total of 34 cysteines distributed throughout the whole sequence (Figure 1A and S1). C185 is among 12 cysteines predicted to have a higher accessible surface area (ASA) (ASA>20%) (Figure 1B).^49^ The AlphaFold structure predicts that C185 is located at the α-helix bundle between PX and LRR, where its thiol group is exposed to the surface and surrounded by charged residues (K217 and D182) (Figure 1A). C185 is predicted to have a similar or reduced pK_a_ value (pK_a_ = 8.8) compared to other cysteines (average pKa = 9.68) (Figure S1A).^50^ C185 and its neighboring sequence are conserved in mammals, and C185 conservation was also found in birds (Gallus gallus) and fish (Danio rerio) (Figure 1C).

**Figure 1.**
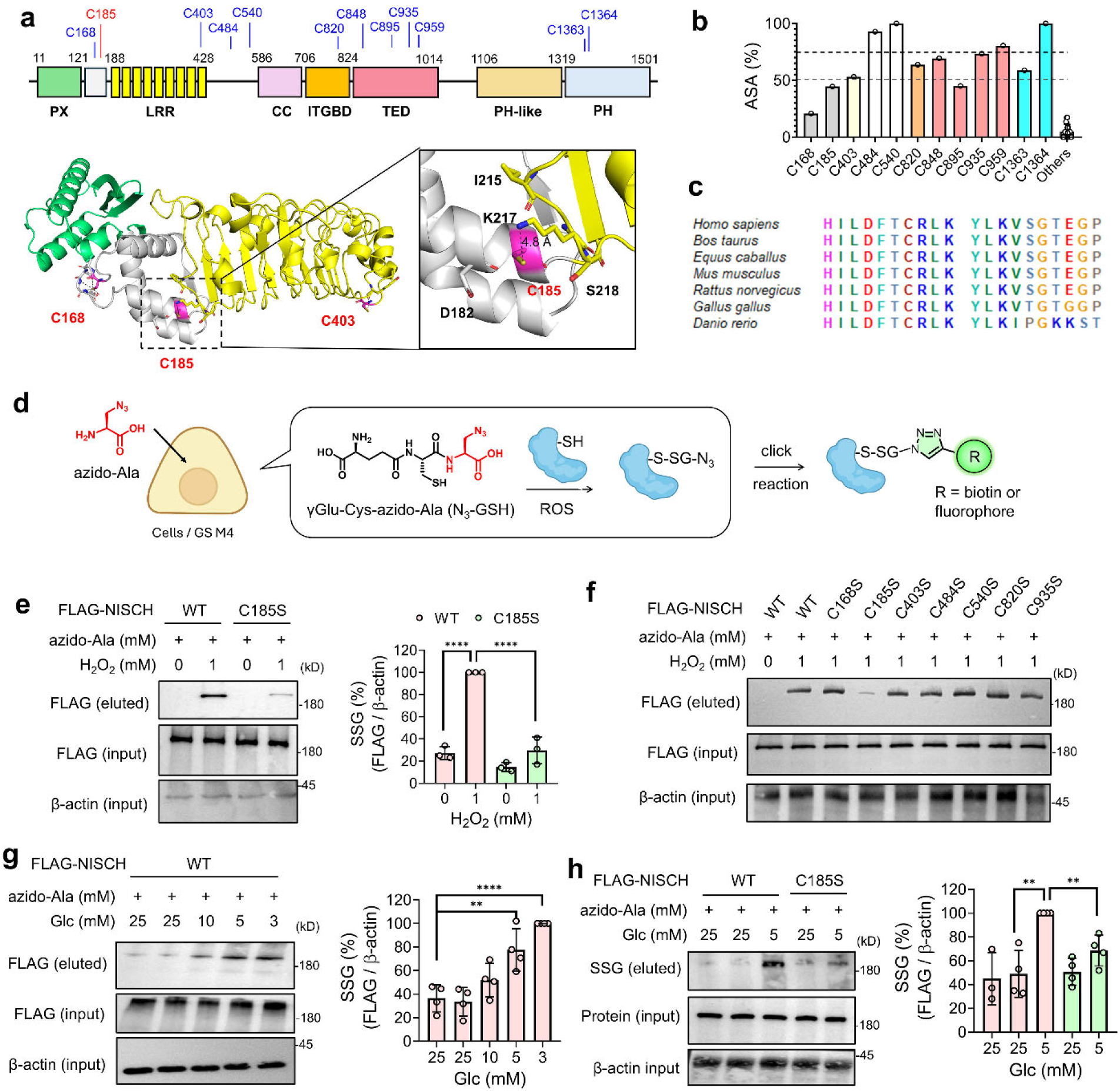
NISCH is selectively susceptible to glutathionylation at C185. (**a**) NISCH structure, domains, and selected cysteines, including C185 (stick model). The structure is retrieved from the AlphaFold database. PX, LRR, CC, and ITGBD were found in UniProt. TED, PH-like, and PH are found in the AlphaFold and Pfam databases. (**b**) The accessible surface area (ASA) of cysteines in NISCH. (**c**) C185 and its neighboring residues in NISCH orthologs. (**d**) The outline of the clickable glutathione approach. Upon incubating azido-Ala, cells expressing a glutathione synthetase mutant (GS M4) synthesize and produce clickable glutathione (N_3_-GSH), which forms glutathionylation with proteins. (**e-f**) Glutathionylation of NISCH WT and cysteine mutants in HEK293/GS M4 cells in response to hydrogen peroxide (H_2_O_2_). H_2_O_2_ was incubated for 15 min (n=3). (**g-h**) Glutathionylation of NISCH WT and C185S in MDA-MB-231 cells expressing GS M4 in low glucose (LG) compared to high glucose (HG) conditions. Cells were incubated in LG or HG for 16 h (n=3-4). The lysates were subjected to click chemistry using biotin-alkyne and analyzed by Western blot analysis before (input) and after (eluted) pull-down with streptavidin-agarose. Data represent the mean ± SD. The statistical difference was analyzed by one-way ANOVA and Tukey’s post-hoc test (**e**, **g**-**h**), where *p < 0.03, **p < 0.002, ***p < 0.0002, ****p < 0.0001.

To demonstrate glutathionylation of NISCH at C185, NISCH WT and C185S were expressed and compared for S-glutathionylation using our clickable glutathione approach.^51,52^ In this approach (Figure 1D), a glutathione synthetase mutant (GS M4) was expressed in cells, which were then incubated with azido-Ala, resulting in the biosynthesis of clickable glutathione (azido-glutathione, N_3_-GSH) in cells. After inducing glutathionylation, the click chemistry-based biotinylation and the subsequent analysis enabled the detection of glutathionylation by azido-glutathione (Figure 1D).^51,52^ First, a bolus of H_2_O_2_ addition in HEK293 cells stably expressing GS M4 (HEK293/GS M4)^53^ induced high levels of S-glutathionylation at the proteome level (Figure S2A). Consistently, H_2_O_2_ addition caused significant S-glutathionylation in NISCH WT (Figure 1E, lane 2 vs. 1). In contrast, H_2_O_2_ caused no or a weak signal of glutathionylation in NISCH C185S (Figure 1E, lane 2 vs. 4), supporting the glutathionylation at C185. To corroborate the selective glutathionylation at C185, we prepared and analyzed six other mutants of cysteines (C168S, C403S, C484S, C540S, C820S, and C935S) that are predicted to have high ASA (Figure 1B). Impressively, in response to H_2_O_2_, all cysteine mutants except C185S showed S-glutathionylation comparable to NISCH WT (Figure 1F and S2B), while C185S showed a dramatically reduced signal of S-glutathionylation. These experiments support that NISCH is selectively glutathionylated at C185. The data are consistent with our previous proteomic analysis for S-glutathionylation in response to H_2_O_2_, where LC-MS/MS analysis identified the mouse NISCH-derived peptide glutathionylated only at C186 (TDLGHILDFTC_186_*R, mouse C186 is equivalent to human C185) (Figure S2C).^54^

Previously, we demonstrated that low glucose (5 and 1 mM) compared to high glucose concentrations (25 mM) increases ROS production and global glutathionylation in various cell lines, including MDA-MB-231 and MCF7 cells,^17,21,27,53^ likely resulting from reduced levels of cellular NADPH synthesis in glucose-addicted cancer cells.^55–57^ Along with ROS production, lower glucose conditions were shown to increase cell migration in MCF7 and MDA-MB-231 cell lines.^17,27^ Lower glucose conditions may represent tissues or solid tumors with higher metabolic activity but with insufficient vasculature,^58,59^ resulting in glucose- or nutrient-starved conditions and inducing cell migration for metabolic adaptation.^60,61^ The clickable glutathione approach demonstrated that lower concentrations of glucose (i.e., 3-5 mM) resulted in high levels of glutathionylation in NISCH WT (Figure 1G, lanes 4-5 vs. 1-3) in MDA-MB-231 cells. Under the same conditions, the glutathionylation levels were significantly reduced in NISCH C185S (Figure 1H, lanes 5 vs. 3, and Figure S2D), further supporting selective NISCH glutathionylation at C185.

### NISCH S-glutathionylation increases cell migration and invasion

To demonstrate the role of NISCH C185 glutathionylation, we used the MCF7 cell line, which expresses a high level of endogenous NISCH (Figure S6A),^30^ and generated the NISCH knockout (KO) cell line using the CRISPR/Cas9 system (Figure 2A and S3A). The endogenous NISCH KO was confirmed in two cell lines (Figure 2A, colony #18 and #16). The in vitro scratch assay showed that both NISCH KO cell lines (#18 and #16) migrate faster than the parental or negative colony MCF7 cell lines that express endogenous NISCH (Figure 2A and S3B), consistent with the negative role of NISCH in cell migration.^29^ Because colonies #18 and #16 showed migration at comparable levels, #18 (named MCF7-NISCH KO or MCF7^NIS-KO^ hereafter) was used in all following assays. To specify the effect of NISCH C185 glutathionylation, NISCH WT or C185S was re-expressed in MCF7^NIS-KO^ cells at comparable levels to endogenous NISCH in MCF7 (Figure S3C). The in vitro scratch assay showed that NISCH WT or C185S re-expression reduced the migration of MCF7^NIS-KO^ cells (Figure 2B, bars 3 and 5 vs. 1), attributed to the inhibitory role of NISCH in cell migration.^29^ However, MCF7^NIS-KO^ cells re-expressing NISCH WT (MCF7^NIS-KO^-NISCH WT) increased cell migration in low glucose (LG, 5 mM) compared to high glucose (HG, 25 mM) (Figure 2B, bars 4 vs. 3), which is not significantly observed in MCF7^NIS-KO^ cells re-expressing NISCH C185S (MCF7^NIS-KO^-NISCH C185S) (Figure 2B, bars 6 vs. 5).

**Figure 2.**
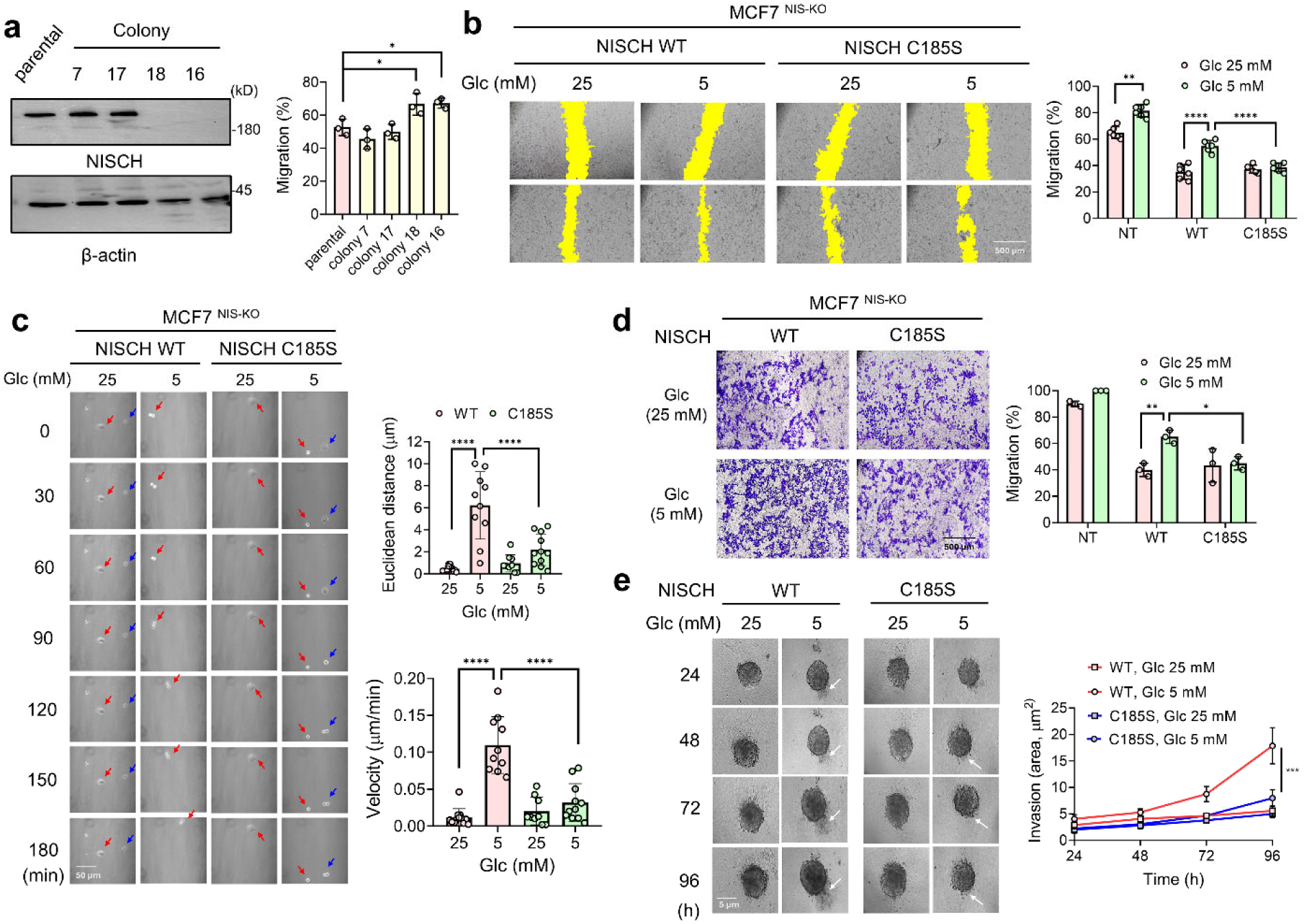
NISCH C185 S-glutathionylation regulates cell migration and invasion. (**a-d**) Migration analysis of MCF7 cells expressing NISCH WT or C185S. (**a**) Analysis of NISCH knockout (KO) in MCF7 cells (left, n=3), and cell migration analysis (right, n=3) after CRISPR/Cas9-based NISCH KO. (**b**) The wound-healing migration analysis. MCF7-NISCH KO (MCF7^NIS-KO^) cells re-expressing NISCH WT or C185S (MCF7-NISCH WT or C185S) were incubated in high glucose (HG, 25 mM) or low glucose (LG, 5 mM) (n=6). The yellow color indicates the area without cells. (**c**) Single-cell migration analysis. MCF7-NISCH WT or C185S cells in LG or HG were monitored by confocal microscopy every 10 min (0-180 min). The images were combined to analyze the migration of individual cells (n=8-10). (**d**) Transwell migration analysis. MCF7-NISCH WT or C185S cells were incubated in LG or HG for 24 h (n=3). (**e**) Spheroid invasion analysis of MCF7-NISCH WT or C185S. Individual spheroids were incubated in HG or LG and monitored daily for 96 h (n=10). Data represent the mean ± SD. The statistical difference was analyzed by one-way ANOVA and Tukey’s post-hoc test (**a**, **c**) or two-way ANOVA and Sidak’s post-hoc test (**b**, **d-e**), where *p < 0.03, **p < 0.002, ***p < 0.0002, ****p < 0.0001.

As opposed to collective cell migration in the scratch assay, two cohorts of cells with WT or C185S were further examined for single-cell migration. Cells expressing NISCH WT or C185S were plated at low confluency, and their migrations were monitored in a confocal microscope over 3 h. MCF7^NIS-KO^-NISCH-WT cells migrated at higher velocities and with longer distances in LG versus HG (Figure 2C and S4). In contrast, MCF7^NIS-KO^-NISCH-C185S cells did not increase velocities and distances of migration significantly in LG compared to HG (Figure 2C and S4). In addition, cell migration was analyzed in the transwell migration assay, supporting the same conclusion: NISCH WT and C185S re-expression reduced the migration of MCF7^NIS-KO^ cells (Figure 2D, bars 3 and 5 vs. 1). MCF7^NIS-KO^-NISCH-WT cells increased migration in LG versus HG (Figure 2D, bars 4 vs. 3), whereas MCF7^NIS-KO^-NISCH-C185S cells did not show a significant increase in cell migration in LG (Figure 2D, bars 6 vs. 5).

In addition to the migration assay, the effect of NISCH S-glutathionylation on cell invasion was analyzed. To monitor cell invasion through the 3-dimensional (3-D) matrix, we analyzed cells expressing WT or C185S in the spheroid invasion assay. MCF7^NIS-KO^-NISCH-WT cells exhibited similar sizes of spheroids in both LG and HG but displayed higher spreading or outgrowth of cells into the matrix in LG compared to HG (Figure 2E and S5). In contrast, MCF7^NIS-KO^-NISCH-C185S cells resulted in less significant outgrowth relative to MCF7^NIS-KO^-NISCH-WT in LG (Figure 2E).

Lastly, to extend the analysis beyond the MCF7 cell line, we investigated similar analyses with MDA-MB-231 cells, which harbor a lower level of endogenous NISCH compared to MCF7 (Figure S6A). NISCH constructs (WT or C185S) were expressed in MDA-MB-231 (ca. 2-fold higher levels than endogenous NISCH) (Figure S6B), which were compared in in vitro migration and transwell invasion assays in LG and HG conditions. The expression of NISCH WT or C185S significantly reduced the migration and invasion of MDA-MB-231 cells (Figure S6C-D). However, NISCH WT-expressing cells increased migration and invasion in LG compared to HG, which was insignificantly observed in NISCH C185S-expressing cells (Figure S6C-D). These experiments demonstrate that NISCH C185 oxidation or S-glutathionylation enhances cancer cell migration and invasion in oxidative stress.

### NISCH S-glutathionylation induces its dissociation from Rac1 and PAK1

NISCH is a negative regulator of cell migration via binding and inhibiting signaling proteins, including integrin α5, PAK1, Rac1, and LIMK1.^30^ Previous studies have reported that N-terminal NISCH domains (LRR, CC, and ITGBD) are the major regions that interact with PAK1 and Rac1 (Figure 3A).^33,34^ NISCH’s N-terminal CC and ITGBD domains also bind to LIMK1,^37^ while NISCH’s ITGBD interacts with integrin α5 (Figure 3A).^30,47^ C185 is located proximally to LRR, which participates in protein-protein interactions. Therefore, we hypothesized that NISCH C185 glutathionylation or oxidation may inhibit protein-protein interactions with PAK1, Rac1, or other proteins, thereby activating them and their downstream pathways to increase cell migration and invasion.

**Figure 3.**
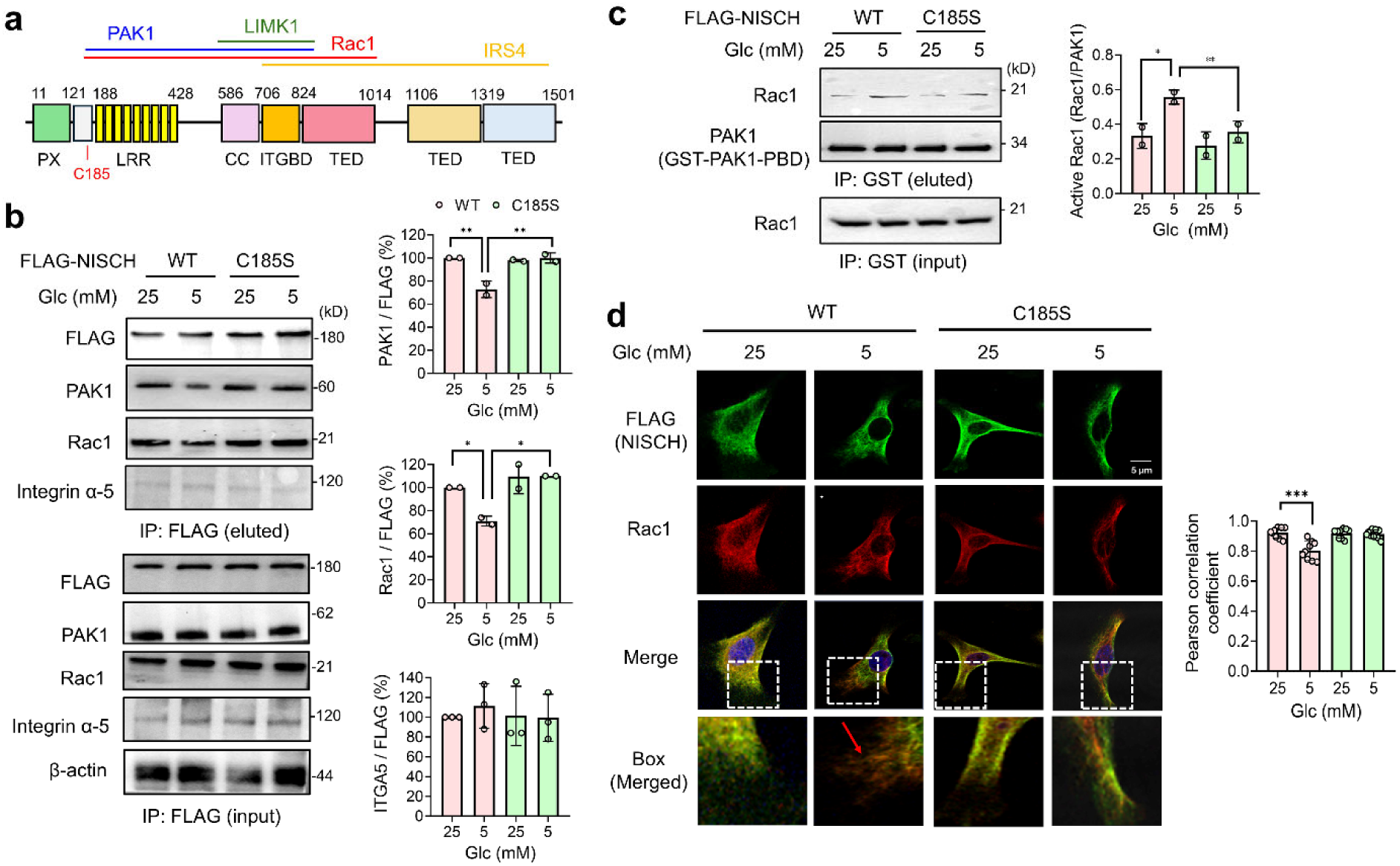
NISCH S-glutathionylation disrupts protein-protein interactions, activating the Rac1 pathway. (**a**) NISCH domain and its interacting partners. NISCH domains interacting with protein partners were drawn in lines in different colors. (**b-d**) Analysis of NISCH-protein interactions and Rac1 downstream pathway in MDA-MB-231 cells. MDA-MB-231 cells expressing NISCH WT or C185S were incubated in low or high glucose (LG or HG). (**b**) NISCH glutathionylation dissociates NISCH from Rac1 and PAK1. After incubating cells in LG or HG for 16 h, the co-immunoprecipitation was used to examine the binding of Rac1, PAK1, or integrin α5 (n=2-3). (**c**) Rac1 activation analysis after NISCH C185 glutathionylation. After incubating cells for 16 h, the GST-fused p21-binding domain (PBD) derived from PAK1 (GST-PAK1-PBD) was used to pull down the Rac1-GTP form, which was then analyzed by Western blot (n=2). (**d**) The co-localization analysis of NISCH and Rac1. After 16 h, cells were fixed and analyzed by antibodies to FLAG (green) and Rac1/Cdc42 (red) (n=10 images). The merged images were analyzed to examine the relative displacement of NISCH and Rac1 in the cell periphery at higher magnification (box). Data represent the mean ± SD. The statistical difference was analyzed by one-way ANOVA and Tukey’s post-hoc test (**b-d**), where *p < 0.03, **p < 0.002, ***p < 0.0002, ****p < 0.0001.

Therefore, we examined NISCH interactions with PAK1 and Rac1 via co-immunoprecipitation (co-IP). In MDA-MB-231 cells, NISCH WT showed reduced binding with PAK1 and Rac1 in LG compared to HG (Figure 3B, lane 2 vs. 1). In contrast, NISCH C185S maintained its binding with PAK1 and Rac1 in LG and HG conditions (Figure 3B, lane 4 vs. 3). Similarly, the proximity ligation assays (PLA) further confirm that NISCH decreases its binding to PAK1 and Rac1 in cells expressing NISCH WT, compared to C185S, in LG (Figure S7). On the other hand, the co-IP experiment indicates that NISCH’s interaction with integrin 5α remained unchanged in cells expressing NISCH WT or C185S in both LG and HG conditions (Figure 3B), which is attributed to the fact that C185 is located distant from the integrin binding region (i.e., ITGBD) in NISCH (Figure 3A). These experiments support that NISCH C185 glutathionylation induces NISCH’s dissociation from PAK1 and Rac1.

### NISCH S-glutathionylation activates Rac1 and its downstream pathway

NISCH-Rac1 dissociation is reported to activate Rac1 and its downstream pathways.^34^ Consistently, the Rac1-GTP pull-down experiment showed a higher amount of active Rac1 in MDA-MB-231 cells expressing NISCH WT in LG versus HG (Figure 3C, lane 2 vs. 1). In contrast, Rac1 was insignificantly activated in cells expressing NISCH C185S (Figure 3C, lane 4 vs. 3), supporting that NISCH C185 glutathionylation activates Rac1 upon NISCH-Rac1 dissociation.

The activated Rac1 moves to the cell periphery,^62^ where it promotes actin polymerization (i.e., lamellipodia formation) via recruiting the WAVE complex and activating Arp2/3 for actin branching, which is further stabilized by cortactin.^63,64^ Therefore, we examined the co-immunostaining of NISCH and Rac1. In MDA-MB-231 cells expressing NISCH WT, both NISCH and Rac1 (or Cdc42 detected via Rac1/Cdc42 antibody) were found to be co-distributed throughout the cytoplasm in HG (Figure 3D, column 1, and bar 1, Figure S8). However, in LG, the signal for Rac1/Cdc42 was higher than NISCH in the cell periphery (Figure 3D, column 2, and bar 2, Figure S8), suggesting an increased localization of Rac1/Cdc42 to the cell membrane. In contrast, the higher localization of Rac1/Cdc42 was not observed in cells expressing NISCH C185S in both LG and HG conditions (Figure 3D, column 3-4, and bar 3-4, Figure S8), supporting the potential translocation of Rac1 (or Cdc42) to the cell membrane upon Rac1 activation, which is implicated in lamellipodia formation for enhanced migration.^63^

### NISCH S-glutathionylation activates PAK1-LIMK1-cofilin pathways

PAK1 regulates multiple signaling pathways to increase cell migration. For example, PAK1 activates the LIMK1-cofilin axis to regulate actin filament dynamics.^36^ PAK1 also activates the upstream MAPK pathway, leading to the activation of ERK.^36^ PAK1 also regulates focal adhesion formation and dynamics by phosphorylating paxillin, an adaptor protein in the focal adhesion complex, recruiting signaling molecules.^65^

To demonstrate the effects of NISCH-PAK1 dissociation resulting from NISCH S-glutathionylation, we analyzed the activation of PAK1 and its downstream pathways. The overexpression of NISCH constructs (WT or C185S) in MDA-MB-231 cells decreased the phosphorylation of PAK1, LIMK1, and cofilin (Figure 4A-B, lane 2 vs. 1), indicating that NISCH negatively regulates their activation states. Importantly, phosphorylation levels of PAK1, LIMK1, and cofilin were increased in MDA-MB-231 cells expressing NISCH WT in LG compared to HG (Figure 4A-B, lane 3 vs. 2), which was not seen in cells expressing NISCH C185S (Figure 4A-B, lane 5 vs. 4), supporting that the PAK1-LIMK1-cofilin axis is activated upon NISCH C185 glutathionylation or its dissociation from NISCH. In addition, phosphorylation levels of ERK1/2 and MEK1/2 were reduced upon overexpression of NISCH WT or C185S (Figure 4A, lane 2 vs. 1), supporting their potential regulation via NISCH. However, their phosphorylation levels were not altered in cells expressing NISCH WT or C185S in LG compared to HG (Figure 4A, lanes 2-5), suggesting that ERK, despite a potential PAK1 downstream target, was not activated after NISCH C185 glutathionylation. Similarly, phosphorylation of paxillin, the PAK1 substrate important for focal adhesion,^65^ remains unchanged in both cells expressing WT or C185 in LG and HG conditions (Figure 4C, lane 2-5), suggesting regulation of selected PAK1 downstream targets after NISCH C185 glutathionylation. These data support our model (Figure 4D) that NISCH retards cell migration and invasion by binding and inhibiting signaling proteins, including Rac1 and PAK1, under non-stressed conditions. However, in response to ROS, glutathionylated NISCH decreases its interaction with Rac1 and PAK1, resulting in activation of the Rac1/PAK1-LIMK1-cofilin pathways, which leads to cytoskeletal actin reorganization and increased cell migration.

**Figure 4.**
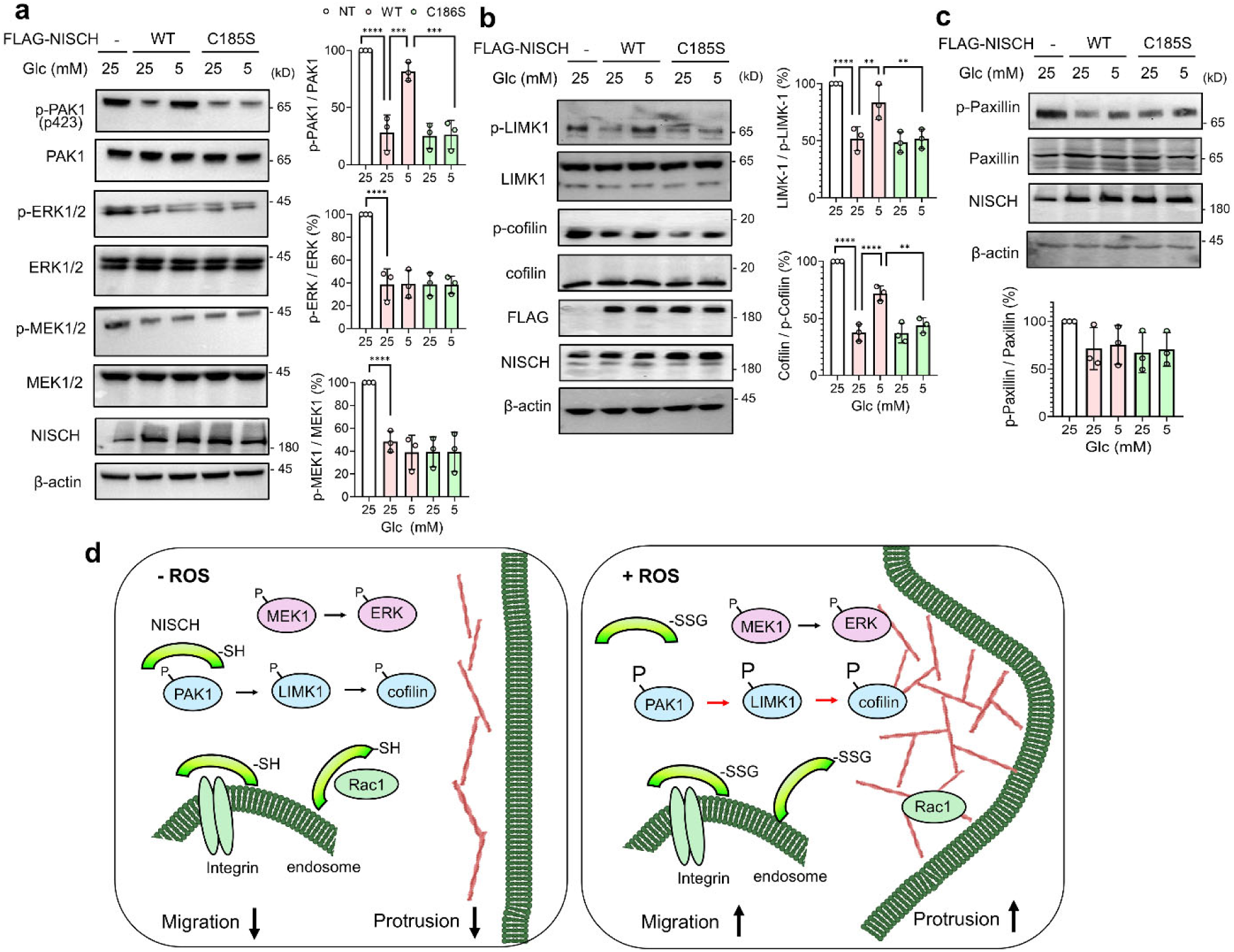
NISCH S-glutathionylation activates the PAK1-LIMK1-cofilin pathway. MDA-MB-231 cells expressing NISCH WT or C185S were incubated in low or high glucose (LG or HG) for 16 h. (**a**) Phosphorylation levels of PAK1, ERK1/2, and MEK1/2 (n=3). (**b**) Phosphorylation levels of LIMK1 and cofilin (n=3). (**c**) Paxillin phosphorylation level (n=3). (**d**) A model for cell migration induced by NISCH glutathionylation. In a non-stressed condition, NISCH binds to its protein partners, including Rac1 and PAK1, thus suppressing Rac1- and PAK1-mediated cell migration. NISCH may bind and retain Rac1 and integrin alpha5 in the cytosol or endosome. However, upon ROS production, NISCH is glutathionylated at C185, dissociating Rac1 and PAK1, but not integrin alpha5. The activated PAK1 activates its downstream LIMK1 and inhibits cofilin through phosphorylation, thereby increasing actin dynamics and polymerization. The dissociated Rac1 increases its localization to the membrane, where it activates actin filament branching, induces lamellipodia and membrane ruffles, and increases cell migration. Data represent the mean ± SD. The statistical difference was analyzed by one-way ANOVA and Tukey’s post-hoc test (**a**-**c**), where *p < 0.03, **p < 0.002, ***p < 0.0002, ****p < 0.0001.

### Engineered NISCH mutant suppresses cancer cell migration and invasion

The Gene Expression Profiling Interactive Analysis (GEPIA)^66^ and the Human Protein Atlas Database^67^ indicate lower RNA and protein expressions of NISCH in all types of breast tumors compared to normal tissues (Figure S9), consistent with NISCH’s role as a tumor suppressor. Our data also suggest that the NISCH C185S construct, unable to form S-glutathionylation, is more effective in suppressing cancer cell migration and invasion than NISCH WT, under oxidative stress conditions (Figure 2B-E, and S6C-D). Therefore, we sought to generate a recombinant NISCH C185S construct and examine the effects of its intracellular delivery on suppressing cancer cell migration and invasion. However, NISCH is a relatively large protein (ca. 166 kDa) with many flexible loops, which makes it challenging to generate a recombinant protein construct. Expectedly, we failed to express and purify the recombinant N-terminal construct of NISCH (amino acids 1-824) in bacterial cultures. Therefore, we used an engineered NISCH construct (amino acids 1-468, encompassing PX and LRR, named NISCH PX-LRR), which was recently developed by introducing stabilizing mutations predicted by the PROSS algorithm (Figure S10A).^68^ Its crystal structure (PDB: 8ESF) closely aligns with the AlphaFold structure (Figure S10B). Considering our data of S-glutathionylation, additional mutations (i.e., C185S, C184T, C298S, and C300E: the last three mutations revert Cys in NISCH PX-LRR to original amino acids in NISCH WT, while the C185S mutation was added to examine C185 glutathionylation) were introduced to generate the second engineered NISCH constructs (named enNISCH WT and C185S, which contain Cys and Ser at the 185 position, respectively) (Figure S10A).

To demonstrate the effects of enNISCH (and its S-glutathionylated form) in regulating cancer cell migration and invasion, we applied our recent G-PROV approach^28^ (Figure 5A). In this approach, a glutathione derivative, dehydroglutathione (dhG), introduces a non-reducible mimic of glutathionylation onto a protein of interest (POI), which is then delivered to cells via a fusogenic liposome (Figure 5A). The effect of POI glutathionylation can then be investigated in these cells.^28^ dhG incubation with purified enNISCH WT or C185S (Figure S11A) induced glutathione modification onto enNISCH WT, detected by glutathione antibody (Figure 5B). In contrast, dhG modification was less significant in enNISCH C185S, indicating selective dhG modification at C185 (Figure 5B) and resembling the pattern of S-glutathionylation on enNISCH (Figure S11B-C). Next, enNISCH constructs (WT and C185S) incubated without and with dhG modification (i.e., enNISCH WT, enNISCH WT-SG, enNISCH C185S, and enNISCH C185S-SG) were individually prepared in a fusogenic liposome, which delivers its cargo mainly into the cytoplasm.^28,69^ The four fusogenic liposomes were then incubated with MDA-MB-231 cells under non-stressed conditions (i.e., 25 mM Glc). Western blot analyses of the lysates confirmed comparable levels of four enNISCH constructs delivered to cells (His blot, Figure 5C, lanes 3-6), with enNISCH WT-SG displaying a significant level of glutathione modification compared to the other constructs (Figure 5C, lane 4 vs. 3 and 5-6). Notably, enNISCH WT-SG is the only protein with apparent glutathione modification in the proteome (lane 4, Figure 5C).

**Figure 5.**
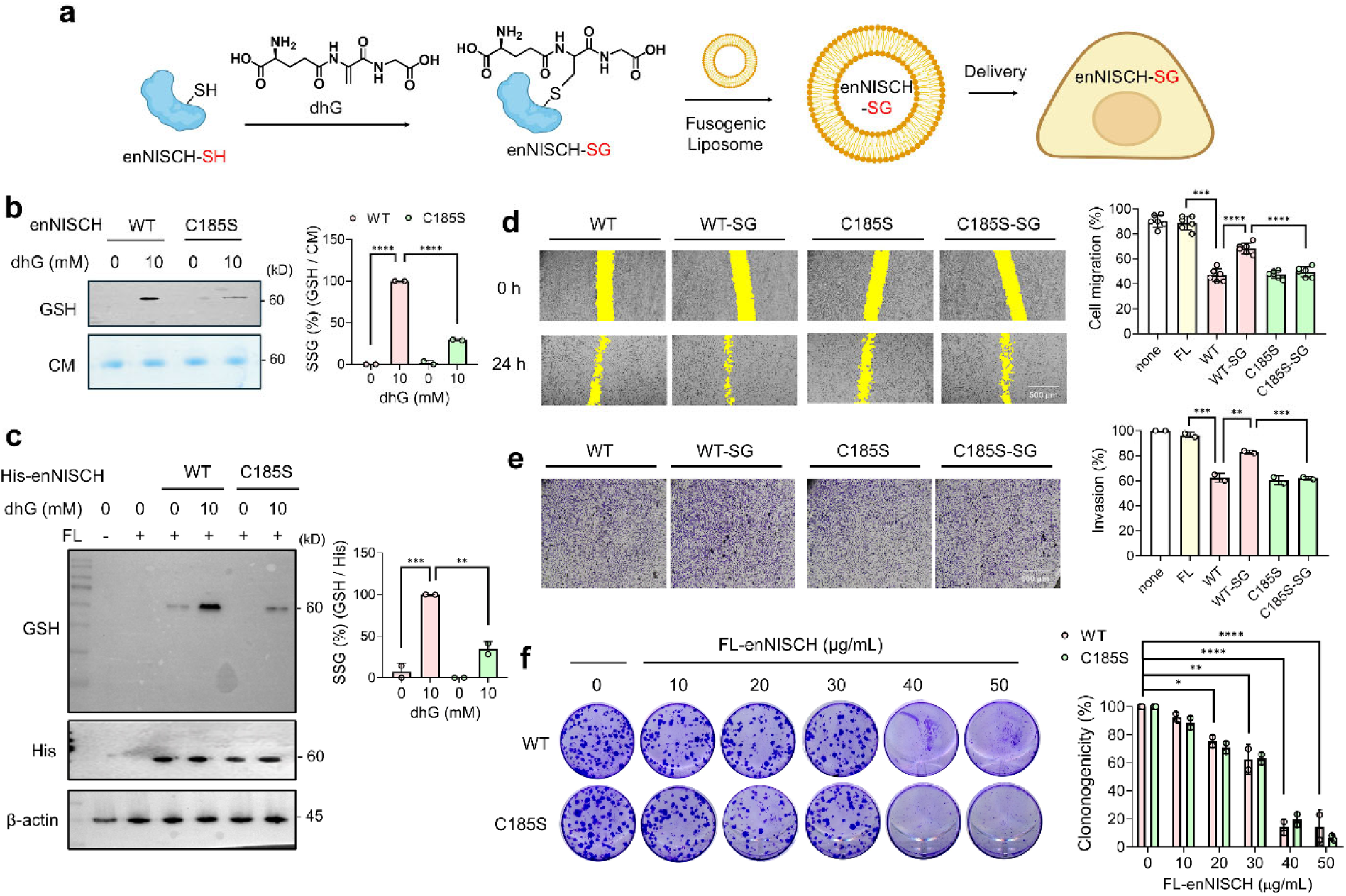
Engineered NISCH mutant suppresses cancer cell migration and invasion. (**a**) A scheme of the G-PROV approach that enables functional analysis of glutathionylation on the protein of interest (POI): dehydroglutathione (dhG) is used to introduce a non-reducible glutathione modification on the purified engineered NISCH construct (enNISCH). Subsequently, the glutathione-modified NISCH construct is delivered to cells via fusogenic liposome, where the effect of glutathionylation on NISCH can be analyzed. (**b**) dhG modification on purified enNISCH. Purified enNISCH WT or C185S were incubated with dhG. The dhG-mediated glutathione modification was analyzed by glutathione antibody (n=2). (**c**) Analysis of enNISCH delivered to MDA-MB-231 cells via fusogenic liposome. After incubating fusogenic liposomes containing His-enNISCH constructs (modified or non-modified by dhG) for 2 h, lysates were analyzed by Western blot (n=2). (**d-e**) The wound-healing migration assay. After incubating fusogenic liposomes for 2 h, MDA-MB-231 cells were analyzed by the wound-healing migration (**d**, n=6) and transwell invasion (**e**, n=2) assays. (**f**) The colony formation by enNISCH constructs. Fusogenic liposomes containing enNISCH WT or C185S were incubated with MDA-MB-231 cells on plates, and the number of colonies formed was counted after 12 days. For **d**-**f**, cells were incubated in DMEM (i.e., 25 mM Glc). The statistical difference was analyzed by one-way ANOVA and Tukey’s post-hoc test (**b**-**e**) or two-way ANOVA and Sidak’s post-hoc test (**f**), where *p < 0.03, **p < 0.002, ***p < 0.0002, ****p < 0.0001.

Encouragingly, the in vitro scratch migration assay demonstrated that the delivery of enNISCH WT or C185S via fusogenic liposome suppressed the migration of MDA-MB-231 cells (Figure 5D, bars 3 and 5 vs. 1, and S11D). Importantly, enNISCH WT-SG caused higher cell migration, compared to three other constructs (enNISCH WT, C185S, and C185S-SG) (Figure 5D, bars 4). Similarly, the transwell invasion assay demonstrated that the delivery of non-modified enNISCH (WT or C185S) suppresses cell invasion (Figure 5E, bars 3 and 5 vs. 1, and S11E). In contrast, dhG-modified enNISCH WT (enNISCH WT-SG) showed higher invasion compared to dhG-treated enNISCH C185S (Figure 5E, bars 4 vs. 6). In biochemical analysis, enNISCH WT-SG showed a modest but significant reduction in binding in vitro with purified Rac1 compared to the other three constructs (Figure S11F). In addition, enNISCH WT-SG delivered to MDA-MB-231 cells caused PAK1 phosphorylation, compared to the other three constructs (Figure S11G). These data support that dhG-mediated glutathione modification in enNISCH resembles the effect of NISCH S-glutathionylation and increases cell migration and invasion. Furthermore, the data suggest that the enNISCH C185S construct, which is devoid of the S-glutathionylation site, could suppress the migration and invasion of MDA-MB-231 cells more effectively than its WT counterpart under oxidative stress.

Lastly, enNISCH constructs (WT and C185S without dhG modification) were examined for their potential in suppressing the colony formation of MDA-MB-231 cells. The delivery of fusogenic liposomes containing enNISCH WT or C185S into the metastatic MDA-MB-231 cells resulted in a lower number of colonies, compared to fusogenic liposome alone (20-50 versus 0 μg/mL, Figure 5F), suggesting that the engineered NISCH construct (both WT and C185S) retains the potential for effective inhibition of clonogenicity and colony formation.

## Discussion

Evidence is emerging to support the significance of ROS in regulating cell migration, adhesion, invasion, and EMT, while contributing to cancer progression and metastasis.^5,6^ Because ROS regulates proteins and enzymes in all major biological processes via cysteine oxidations, including S-glutathionylation, the primary questions are the identification and mechanistic analysis of proteins and enzymes that are regulated via cysteine oxidations.^13^ In this perspective, examples of cysteine oxidations in regulating cell migration, invasion, and EMT have been reported over the years. For example, MAP kinase phosphatase 1 (MKP-1) in monocytes and macrophages is glutathionylated in response to metabolic stress,^22^ thereby enhancing migration and adhesion, which promotes plaque formation and atherosclerosis.^70^ Metabolic stress also induces glutathionylation and degradation of 14-3-3 zeta, resulting in hypermigratory and proatherogenic monocytes.^26^ NADPH oxidase-induced ROS causes glutathionylation of actin, increasing actin dynamics and controlling chemotaxis in neutrophils.^19^ S-glutathionylation of LMW-PTP increased endothelial cell migration induced by VEGF.^25^ We also reported that Ser/Thr phosphatase PP2Cα is glutathionylated in response to EGF, thereby increasing epithelial cell migration and invasion.^17^ Glutathionylation of p120-catenin, a key regulator of E-cadherin in cell-cell adhesion, induces E-cadherin degradation, increasing epithelial cell migration and invasion.^27^ Fatty acid binding protein 5 (FABP5) glutathionylation enhances its binding affinity for fatty acids, such as linoleic acid, facilitating its translocation to the nucleus, activation of PPARβ/δ, and increased cell migration.^28^ Beyond S-glutathionylation, EGF-induced ROS causes EGFR cysteine sulfenylation, enhancing the kinase activity associated with increased cell migration and proliferation.^71^ In addition, motile cells generate H_2_O_2_ at the protrusions, inducing cofilin oxidation that contributes to cell polarization and migration.^8^ 15-PGDH cysteine is oxidized to sulfonic acid, which causes its degradation and the EMT process.^72^ Furthermore, the global level of cysteine oxidation induced by EGF^73^ increases the expectation of discovering additional redox-dependent proteins and cysteines that regulate cell migration, adhesion, and invasion.

In this report, we demonstrate that NISCH contains the redox-regulatory cysteine (i.e., Cys185), and its glutathionylation enhances cell migration and invasion. First, it is interesting to find that NISCH has 34 cysteines, among which C185 is selective for S-glutathionylation, although other cysteines are predicted to have surface exposures (i.e., ASA) and pK_a_ values comparable to C185, two factors contributing to the cysteine’s oxidation susceptibility.^74^ However, it has been argued that other factors beyond pK_a_ and ASA contribute to the cysteine oxidation susceptibility in cells,^75^ including protein conformation and protein-protein interactions,^13,27^ which may explain the selectivity for C185 in NISCH. Second, NISCH has been extensively studied in breast cancer. Therefore, we demonstrated migration and invasion of two breast cancer cell lines, MCF7 and MDA-MB-231, upon NISCH S-glutathionylation. However, NISCH is ubiquitously expressed in all major organs and tissues, including neurons.^29^ Therefore, NISCH S-glutathionylation may be important for migration and invasion of other cell types, such as neuron migration and neurite outgrowth.^42^

Mechanistically, we found that NISCH C185 glutathionylation induces the dissociation of NISCH from Rac1 and PAK1, activating Rac1 and PAK1-LIMK1-cofilin pathways that hinge on actin branching and filament formation for the increased protrusion (i.e., lamellipodia). At a molecular level, Cys185 is located in the alpha-helix bundle, proximal to the LRR domain, which is involved in protein-protein interactions. Therefore, our data suggest that the alpha-helix bundle and LRR surrounding Cys185 may serve as direct interaction regions with Rac1 and PAK1 or regulate NISCH conformations that disrupt interactions with Rac1 and PAK1, which is in agreement with previous reports.^33,34^ In addition to Rac1 and PAK1, NISCH is known to interact with other partners, including integrin α5,^31^ the insulin receptor,^76^ and LIMK1.^37^ Our data showed that NISCH C185 glutathionylation does not disrupt the interaction with integrin α5, which is in line with the fact that the integrin binding domain (amino acids 706-824) in NISCH is distant from C185. The insulin receptor is reported to bind the C-terminal domain of NISCH.^30,76^ Thus, it is unlikely to be dissociated from NISCH upon C185 glutathionylation. LIMK1 binds to the N-terminal domains (CC and ITGBD: amino acids 586-824) of NISCH.^47^ Although we did not examine the direct interaction of NISCH with LIMK1 upon NISCH glutathionylation, we showed that LIMK1 is phosphorylated and activated after NISCH C185 glutathionylation, which may result from the dissociation of LIMK1 from NISCH or be attributed to activated PAK1, an upstream kinase of LIMK1. Lastly, the PX domain in NISCH is reported to interact with phosphoinositide and target NISCH to the endosome.^77^ However, our immunostaining was unable to resolve any apparent change in NISCH localization. Additional experiments will be necessary to fully understand the molecular events upon NISCH S-glutathionylation.

Lastly, because of the inhibitory effects of NISCH C185S over WT in cell migration and invasion, we sought to test the therapeutic potential of purified NISCH C185S in suppressing the metastatic potential of the aggressive breast cancer cell line, MDA-MB-231. Unfortunately, NISCH full-length or N-terminal domains (amino acids 1-824 or 1-428) were not soluble after expression in bacterial cells. We tested the recently developed engineered NISCH (amino acids 1-428),^68^ which was expressed and purified in high yield. The reported X-ray crystal structure^68^ shows the folded structures of PX and LRR domains. Interestingly, we demonstrated that enNISCH binds to Rac1 in vitro and suppresses PAK1 phosphorylation in cells, whereas enNISCH C185 glutathionylation causes Rac1 dissociation and elevates PAK1 phosphorylation, further supporting that NISCH PX-LRR domains are important for binding to Rac1 and PAK1. Based on these findings, enNISCH constructs were delivered to MDA-MB-231 cells via fusogenic liposomes. Importantly, enNISCH constructs demonstrated the inhibitory potential in the migration, invasion, and colony formation of MDA-MB-231 cells. Future studies may develop and evaluate smaller versions of the NISCH constructs or apply cancer-targeting liposomes to assess the therapeutic potential of enNISCH in vivo.

## Materials and Methods

### Cell culture

Human embryonic kidney 293 (HEK293) cell line stably expressing GS M4 (HEK293-GS M4), breast cancer cell line MDA-MB-231 (ATCC, HTB-26), and breast cancer cell line MCF-7 (ATCC, HTB-22) were maintained in Dulbecco’s modified Eagle’s medium (DMEM) supplemented with 10% fetal bovine serum (Hyclone, Cytiva), penicillin (100 units/mL) and streptomycin (100 µg/mL) (Pen-Strep, Invitrogen) at 37°C in a 5% CO_2_ environment.

### Cloning

Full-length Nischarin (Origene, pCMV6-myc-DDK-Nischarin, Cat# RC206026) was used to generate mutants by site-directed mutagenesis. Briefly, forward and reverse primers (Table S1) were used for polymerase chain reactions (PCR). Following amplification, 1 µL of DpnI (10 units) was added directly to the PCR reaction and incubated at 37°C for 1 h. Approximately 5 µL of the DpnI-treated reaction was used to transform 50 µL of competent DH5α cells by heat shock, following standard protocols. Transformants were recovered in SOC medium for 1 h at 37 °C with shaking, plated on kanamycin (50 μg/mL) plates, and grown overnight at 37 °C. The next day, colonies were picked up and grown in LB medium containing kanamycin (50 μg/mL), and plasmid DNA was isolated using the miniprep purification kit. Similarly, enNISCH constructs (pET28a-enNISCH) were prepared by site-directed mutagenesis. pET28a-NISCH-N-terminal domain construct (pET28a-NISCH PX-LRR) plasmid was reported previously.^68^ NISCH PX-LRR plasmid was used with the forward and reverse primers (Table S1) for the site-directed mutagenesis to generate pET28a-enNISCH constructs. The mutagenesis was confirmed by the Sanger sequencing method.

### Transfection

Lipofectamine 3000 (ThermoFisher, Cat# L3000015) transfection was performed when the cells were 70% confluent. Briefly, in an Eppendorf tube, 10 µg plasmid NISCH WT or C185S and 28 µL P3000 were mixed in 500 µL Opti-MEM medium. In a separate tube, 21 µL Lipofectamine-3000 reagent was mixed with 500 µL Opti-MEM medium. The diluted DNA/P3000 complex was mixed with Lipofectamine 3000 in a tube and incubated for 30 min at room temperature. The DNA/Lipofectamine mixture was added to 70% confluent HEK293 cells stably expressing GS M4 (HEK293/GSM4)^53^ in a 10 cm dish and incubated in a humidified incubator at 37°C and 5% CO_2_ for 6 h. The Opti-MEM was replaced by DMEM complete medium, and cells were returned to the humidified incubator at 37°C and 5% CO_2_. For MDA-MB-231 and MCF-7 NISCH-KO cell lines, 10 µg DNA, 28 µL P3000, and 21 µL Lipofectamine-3000 were used for a 10 cm dish.

### Glutathionylation analysis

HEK293/GS M4 cells or MDA-MB-231 cells were transfected with the FLAG-NISCH WT or C185S plasmid. After overnight, GS M4 was expressed in MDA-MB-231 cells by infecting the adenovirus carrying GS M4 (Ad-GS M4, VectorBioLab). Briefly, 7.5 µL of adenovirus (9 x 10^10^ PFU/mL) and 8 µL polybrene (10 mg/mL) were mixed in 500 µL DMEM supplemented with 2% FBS. Cells were incubated with the adenovirus solution for 6 h and then in DMEM with 10% FBS for 18 h. Next, GS M4 expressing cells were incubated with 0.6 mM azido-Ala for 20 h for the synthesis of clickable azido-glutathione. HEK293/GS M4 cells were then treated with 1 mM H_2_O_2_ for 15 min to induce glutathionylation. Alternatively, MDA-MB-231 cells were incubated with glucose-free DMEM medium (Gibco, Cat# 11966025) supplemented with 25 mM or 5 mM glucose, along with 0.3 mM azido-Ala for 24 h.

After induction of glutathionylation, cells were lysed using the RIPA lysis buffer containing protease inhibitor cocktail and 50 mM N-ethylmaleimide (NEM). 1 mg lysate was then precipitated by adding ice-cold acetone (4x volume) and incubated at -20°C for 30 min. After centrifugation at 6000 rpm for 5 min, the supernatant was removed, and proteins were resuspended in a 40 μL buffer containing 1x PBS (pH 7.4), 10x SDS (1 μL), and sonicated until proteins were completely dissolved. The click solution (10 μL) was prepared by mixing 20 mM THPTA (5 μL) in water and 20 mM Cu(I)Br dissolved in DMSO/tBuOH (3:1, v/v, 5 μL), which was added to the protein solution. To the solution mixture, 5 mM TAMRA-alkyne (0.5 μL) or 5 mM biotin-alkyne (0.5 μL) was added. The mixture was incubated for 1 h at room temperature in the dark. For in gel fluorescence analysis, the reaction using TAMRA-alkyne was terminated by adding SDS loading buffer and boiling for 10 min at 95°C. The resulting proteins were resolved on an SDS-PAGE gel, and fluorescent images were captured using an iBright imaging system (Thermo Scientific), and protein level was visualized by Coomassie staining. For monitoring NISCH glutathionylation, the click reaction using biotin-alkyne was precipitated by adding ice-cold acetone (4x volume) and incubating at -20°C for 30 min. After centrifugation at 6000 rpm for 5 min, the supernatant was removed, and proteins were resuspended in a 40 μL buffer containing 1x PBS (pH 7.4) and 10x SDS (1 μL). The resuspended solution was added to prewashed streptavidin-agarose beads (20 μL, Pierce) and incubated overnight at 4 °C. Beads were washed with 1X PBS (3 x 500 μL). Proteins on beads were eluted by adding 50 μL SDS-loading dye (2x) containing β-mercaptoethanol (3 μL) and heating at 95 °C for 10 min. Eluted proteins were separated by SDS-PAGE and transferred to the PVDF membrane. The membrane was blocked with 5% BSA in TBST (50 mM Tris-HCl, 150 mM NaCl, and 0.1% Tween-20) and incubated with a primary antibody solution, containing FLAG (1:1000) or β-actin (1:500), diluted in a blocking buffer overnight at 4°C. The corresponding horseradish peroxidase (HRP)-conjugated secondary antibodies, anti-mouse IgG (1:1000) and anti-rabbit IgG (1:1000), were used to visualize the proteins by chemiluminescence (Super Signal West Pico). Blots were analyzed using the iBright imaging system (Thermo Scientific).

For purified enNISCH proteins, 100 μg purified enNISCH WT and C185S were individually incubated with 2 μL of 25 mM azido-glutathione (N_3_-GSH), and glutathionylation reaction was induced by adding 0.5 mM H_2_O_2_ for 15 min at room temperature. Cysteines were blocked by adding 200 mM iodoacetamide (IAM) and 10% SDS to the reaction mixture and incubating for 20 min in the dark. The reaction mixture was precipitated by adding prechilled acetone and keeping it at -20°C for 30 min. The precipitated protein was centrifuged at 3,500 rpm for 5 min, and the pellet was resuspended in PBS containing 0.1% SDS. The proteins were resolved on SDS-PAGE gel, and fluorescent images were captured using an iBright imaging system (Thermo Scientific), and protein level was visualized by Coomassie staining.

### Western blot

To analyze phosphorylation levels, MDA-MB-231 cells transfected with NISCH WT or C185S were incubated in glucose-free DMEM medium supplemented with 25 mM or 5 mM glucose for 24 h. Cells were then lysed with 200 μL RIPA lysis buffer supplemented with protease inhibitor cocktail (ThermoFisher, Cat# A32955) and PhosphoSTOP phosphatase inhibitor (Roche, Cat# 4906845001). Protein concentration was determined using Bradford assay. About 40 μg protein lysates were resolved by SDS-PAGE and transferred to PVDF membrane. The membrane was blocked with 5% BSA in TBST containing 50 mM Tris-HCl, 150 mM NaCl, and 0.1% Tween-20, and incubated with primary antibody solution, containing phospho-PAK1 (p423) (1:1000; Cell Signaling, Cat# 2601S), PAK1 (1:1000; Cell Signaling, Cat# 2608S), phospho-cofilin-Ser-3 (Cell Signaling, Cat# 3311S) Cofilin (Cell Signaling, Cat# 3318S), phospho-LIMK-1 (Cell Signaling, Cat# 3841S), LIMK1 (Cell Signaling, Cat# 3842S), phospho-paxillin (Cell Signaling, Cat# 441026G), Paxillin (Cell Signaling, Cat# 2542S) NISCH(D6T4X) (Cell Signaling, Cat# 85124S), phospho-p38 MAPK (1:1000; Cell Signaling, Cat# 9211S), p38 MAPK (1:1000; Cell Signaling, Cat# 9212S), phospho-MEK1/2 (1:1000; Cell Signaling, Cat# 9154S), MEK1/2 (1:1000; Cell Signaling, Cat# 9122S), phospho-p44/42 MAPK (ERK1/2) (1:1000; Cell Signaling, Cat# 9101S), p44/42 MAPK (ERK1/2) (1:1000; Cell Signaling, Cat# 9102S), Integrin α-5 (Cell Signaling, Cat# 4705S) or FLAG antibody (1:1000; Millipore-Sigma, Cat# F3165), diluted in a blocking buffer and incubated for overnight at 4°C. The corresponding horseradish peroxidase (HRP)-conjugated secondary antibodies, anti-mouse IgG (1:1000) and anti-rabbit IgG (1:1000), were used to visualize the proteins by chemiluminescence (Super Signal West Pico). Blots were analyzed using the iBright imaging system (Thermo Scientific).

### Co-immunoprecipitation

For co-IP of NISCH and its interacting proteins (PAK1, Rac1, and ITGA5), MDA-MB-231 cells were transfected with FLAG-NISCH-WT or C185S plasmids. After 24 h, cells were incubated in 25 mM or 5 mM glucose in glucose-free DMEM for 24 h. Cells were then lysed with lysis buffer (Tris-pH 8, 150 mM NaCl, 1% NP-40) supplemented with protease inhibitor cocktail (1X) and phospho-STOP. 1 mg of lysate was incubated with 4 μL of FLAG antibody (1:1000; Milipore-Sigma, Cat# F3165) for 1 h at 4 °C. The solution was mixed with prewashed protein G agarose beads (25 µL beads) and incubated overnight at 4°C under constant rotation. Beads were washed three times with 0.5 mL of washing buffer (50 mM Tris-HCl, 150 mM NaCl, and 0.1% Tween-20) for 10 min each. Bound proteins on beads were eluted by adding 50 μL SDS-loading dye (containing 1 mM β-mercaptoethanol) and boiled at 95°C for 10 min. Eluted proteins were resolved by SDS-PAGE and transferred to PVDF membrane. The membrane was blocked with 5% BSA in TBST (50 mM Tris-HCl, 150 mM NaCl, and 0.1% Tween-20) and incubated with primary antibody solutions, such as PAK1 (1:1000), Rac1/Cdc42 (1:1000; Cell Signaling, Cat# 4651S), Igα-5 (1:1000; Cell Signaling, Cat# 4705S), or FLAG antibody (1:1000; Milipore-Sigma, Cat# F3165), diluted in a blocking buffer overnight at 4°C. Appropriate horseradish peroxidase (HRP)-conjugated secondary antibodies were used to visualize the proteins by chemiluminescence. Blots were analyzed using the iBright imaging system (Thermo Scientific).

### Rac1 activation assay

Rac1 activity was determined using a GST-tagged p21-binding domain (PBD) of PAK1 (i.e., GST-PAK1-PBD, amino acids 70-117), which specifically interacts with the GTP-bound (active) form of Rac1. Briefly, MDA-MB-231 cells transfected with NISCH WT or C185S were incubated in glucose-free medium supplemented with 25 mM or 5 mM glucose for 24 h. Cells were washed with ice-cold PBS before lysis in ice-cold Mg²⁺ lysis buffer (25 mM Tris-HCl, pH 7.5, 150 mM NaCl, 5 mM MgCl₂, 1% NP-40, 10% glycerol) supplemented with protease and phosphatase inhibitor cocktails. Lysates were clarified by centrifugation at 14,000 × g for 5 min at 4°C, and equal amounts of lysate (1 mg) were incubated with 100 µg of GST–PAK1-PBD fusion protein pre-bound to glutathione–Sepharose beads for 45 min at 4°C with gentle rotation. Beads were subsequently washed three times with 1X PBS, and bound proteins were eluted by boiling in 2× Laemmli buffer. Samples were resolved by SDS-PAGE and transferred to PVDF membranes, followed by immunoblotting with anti-Rac1 (Cell Signaling Cat# 4651S) and anti-GST (for GST-PAK1-PBD) (1:1000) antibodies to detect the active GTP-bound Rac1 fraction. In parallel, total Rac1 levels were determined from aliquots of the whole-cell lysates. Rac1 activation was expressed as the ratio of GTP-bound Rac1 to total Rac1.

### Migration assay

Non-transfected or transfected cells with NISCH WT or C185S were seeded into a 12-well plate to produce a fully confluent monolayer (1.5 X 10^5^ cells). The next day, cells were serum-starved for 3 h with DMEM. The wound was created using a 10 μL pipette tip, and the detached cells were removed using PBS(1X). Glucose-free DMEM supplemented with 25- or 5-mM glucose was added to the cells. The wound area was imaged at 0 h and 24 h using a light microscope connected to a camera. Images were analyzed using an MRI wound healing tool in ImageJ software.

### Transwell assay

MDA-MB-231 or MCF-7 NISCH KO cells (3 × 10^5^ cells/well), non-transfected or transfected with NISCH WT or C185S, were seeded in the upper chamber coated with Matrigel (for invasion analysis) or without Matrigel (for migration analysis). The upper chamber was filled with 200 μL glucose-free DMEM including 5- or 25-mM glucose. The lower chamber was filled with 700 μL glucose-free DMEM containing 10% FBS. After 24 h, the medium was removed from the lower and upper chambers. Cells were washed 1X PBS (3 x 5 min), and non-invasive cells were removed using a cotton swab. The bottom side of the transwell insert membrane was incubated with 500 μL 4% formaldehyde for 10 min at room temperature, followed by permeabilization with permeabilization buffer (PBS pH 7.4, 0.1% Triton-X100) for 10 min. Inserts were washed with PBS (3 x 5 min) and stained with 500 μL 0.2% crystal violet solution for 20 min at room temperature. Inserts were then extensively washed with 1X PBS (3 x 5 min) to remove all excess dye. Images were captured using a camera attached to a microscope. The stained dye was extracted with 300 µL 33% acetic acid, and absorbance was measured at 560 nm using a Synergy H1 microplate reader (BioTek).

### Immunofluorescence assay

MDA-MB-231 cells transfected with NISCH WT or C185S were seeded in a 35 mm glass-bottom dish coated with fibronectin (0.1%). After incubating in 5 mM or 25 mM glucose conditions, cells were fixed with 4% paraformaldehyde (500 μL) at room temperature for 10 min. After fixation, cells were washed with 1X PBS three times and permeabilized by incubating with permeabilization buffer (0.1% Triton X-100 in 1X PBS) for 10 min at room temperature. Cells were washed with 1X PBS three times and blocked with blocking buffer (0.25% BSA in PBST) for 30 min at room temperature. After blocking, cells were incubated with the primary antibody in blocking buffer (300 μL) overnight at 4 °C. Primary antibodies include rabbit FLAG (1:500; Thermo Fischer, Cat# PA1-984B) and mouse Rac1 antibody (1:500; Thermo Fischer, clone 23A8 #05-389). The next day, the antibody solution was removed and washed with 1X PBS three times and incubated with secondary antibodies (anti-mouse Alexa Fluor 647 (1:1000; Invitrogen, Cat# A-21235) or anti-rabbit Alexa Fluor 555 (1:500; Invitrogen, Cat# A27039) for 1 h at room temperature. Cells were washed three times with 1X PBS and then stained with Prolong Gold Antifade with DAPI (Invitrogen, Cat #P36931) for 10 min at room temperature. Images were captured from a confocal microscope (Zeiss LSM 700) and analyzed using ImageJ. Pearson’s correlation coefficients were calculated by Just Another Colocalization Plugin (JACoP) in ImageJ software.

### Preparation for NISCH knockout cell line

To prepare the plasmid for NISCH knockout using CRISPR/Ca9, the pSpCas9(bb)-2a-puro (px459) plasmid was obtained from Addgene (Cat# 48139). The target region of the gene was analyzed by ENSEMBL (https://useast.ensembl.org/index.html). Using CRISPOR (http://crispor.tefor.net/), sgRNAs were designed (Table S1). The sense and antisense primers were annealed by mixing 1 µL of each primer (100 µM), 1 µL of 10x T4 ligase buffer, 1 µL of T4 PNK, and 6 µL of sterile water. The solution was incubated at 37°C for 30 min. The solution was then heated to 95°C for 5 min and cooled to room temperature. The annealed primer mixture was diluted to 1:200. The pSpCas9(bb)-2a-puro plasmid was digested by the BbSI restriction enzyme. The diluted annealed primers and the linear vector were ligated using T4 DNA ligase to produce pSpCas9(BB)-2A-Puro gRNA1, pSpCas9(BB)-2A-Puro-gRNA2, and pSpCas9(BB)-2A-Puro-gRNA3.

MCF-7 cells were transfected with pSp-Cas9(BB)-2A-gRNA constructs with Lipofectamine 3000. 14 µg of each pSP-Cas9 (BB)-2A-sgRNA (gRNA1, gRNA2, and gRNA3) was mixed with 21 µL Lipofectamine and 28 µL P3000 reagent. After 48 h, cells were incubated in DMEM with 10% FBS, containing puromycin (0.5 μg/mL). After growing for one week, cells were detached by trypsin, diluted to 5 cells/mL, and transferred to a 96-well plate containing 100 µL DMEM with 10% FBS. The single colonies developed over 3-4 weeks were transferred to 12-well plates. When cells were confluent, they were transferred to the 6 cm dish. After the 6 cm dish became confluent, cell stocks were prepared. Lysates were collected and subjected to western blotting with NISCH primary antibody (1:1000).

### Single-cell tracking assay

MCF7-NISCH KO cells were transfected with NISCH WT or C185S plasmid. Cells were incubated with fresh DMEM medium containing G418 (100 μg/mL) for 7 days to select transfected cells. Cells were seeded to produce 10-20% confluency in a glass-bottom 35 mm dish coated with 0.1% fibronectin (MatTek, Cat# P35G-1.5-14-C). Cells were incubated overnight at 37°C with 5% CO_2_. The next day, cells were incubated with 25 mM or 5 mM glucose in glucose-free DMEM. Images were captured every 10 min for 3 h using a 32X phase contrast confocal microscope (Zeiss LSM700). Manual tracking and chemotaxis tools in ImageJ were used to analyze the images. Briefly, time-lapse image stacks were opened in ImageJ, and individual cells were manually tracked using *Plugins/Tracking/Manual Tracking*. For each cell, a new track was initiated, and the cell centroid was marked sequentially across each frame. Once tracking of a cell was completed, a new track was initiated for the next cell. Upon completion of tracking, data were exported in CSV format, which included time intervals, X and Y coordinates, incremental displacement between frames, and cumulative distance migrated. The exported data were subsequently analyzed to calculate total migration distance, average displacement, and migration velocity.

### Spheroid invasion assay

Spheroids were generated using MCF-7 KO cells expressing NISCH WT or C185S by the hanging drop method.^78^ Briefly, after transfection of NISCH WT or C185S, cells expressing NISCH were selected by incubating with G418 (100 μg/mL) for 7 days. Cells were then detached with 0.05% trypsin–EDTA, neutralized, and counted to prepare a single-cell suspension at 2 X 10⁴ cells/mL. Drops of 25–30 µL containing 1500 cells/drop were dispensed onto the inner surface of sterile 100-mm Petri dish lids, which were inverted over base dishes containing sterile PBS to maintain humidity and prevent evaporation. Cultures were incubated undisturbed for 72 h, during which compact spherical aggregates formed.

For spheroid invasion assays, 96-well plates were precoated with Matrigel (30 µL/well) and allowed to solidify for 30 min at room temperature. Preformed spheroids generated by the hanging drop method were then individually transferred into the precoated wells and incubated for 10 min to facilitate attachment. A mixture of Matrigel and collagen I (1:1 ratio) supplemented with DMEM was then overlaid on top of each spheroid and allowed to polymerize for 30 min at 37°C in a CO₂ incubator. After polymerization, DMEM containing high glucose was added to each well, and cultures were incubated for 24 h. At this point, the medium was replaced with 5 mM or 25 mM glucose in glucose-free DMEM, and spheroid invasion was monitored over time. Images were acquired every 24 h using an inverted phase-contrast microscope, and invasion was quantified using ImageJ by measuring changes in spheroid size and outgrowth area relative to baseline. Briefly, the freehand line tool in ImageJ was used to draw a line along the length of the reference object. The pixel length was determined using the Analyze/Measure function. The scale was then set using Analyze/Set Scale, where the measured pixel distance was entered as “Distance in pixels”. The known distance was set to 1000 (µm), and “Unit of length” was specified as µm. The “Global” option was selected to apply the calibration across images. Distances were subsequently measured using the Analyze/Measure function.

### Recombinant protein purification

pET28a-enNISCH plasmids were transformed into BL21 (DE3) competent cells for expression. A single colony was inoculated into 5 mL Luria Broth (LB) medium containing 50 μg/mL kanamycin and grown at 37 °C for 16 h. 5 mL overnight culture was inoculated into 1 L LB medium containing 50 μg/mL kanamycin and grown at 37 °C until OD_600_ reached 0.6. Protein expression was induced with 0.5 mM isopropyl β-D-1-thiogalactopyranoside (IPTG) and incubated at 18°C for 18 h. Cells were pelleted by centrifugation at 5,000 rpm and washed with ice-cold Tris-HCl buffer (pH 7.4). Cells were lysed using a French press with lysis buffer (25 mM Tris, pH 7.4, 300 mM NaCl, 1 mM DTT, and Pierce protease inhibitor, pH 7.4) and subjected to centrifugation at 15,000 rpm at 4°C. The supernatant was transferred to a fresh tube containing pre-washed Ni-NTA agarose beads at 4°C for 2 h. Proteins were eluted using elution buffer (25 mM Tris, 75 mM NaCl, 300 mM imidazole, 1 mM DTT, pH 7.4). Pure fractions were combined and dialyzed in dialysis buffer (25 mM Tris, pH 7.4, 75 mM sodium chloride, 1 mM DTT, and 5% glycerol) overnight at 4°C. GST-PAK1-PBD protein was expressed and purified in similar method. Briefly, The pGEX-PAK1 PBD plasmid (Addgene, Cat. #12217) was used for expression in BL21 (DE3), and the GST-PAK1-PBD was purified using glutathione Sepharose 4B beads. The protein concentration was measured by using the Bradford assay.

### Recombinant enNISCH glutathionylation

Purified enNISCH protein (0.5 mg) was dissolved in 0.5 mL 1X PBS (pH 8.0) and incubated with 10 mM dehydroglutathione (dhG) or 1 mM oxidized glutathione (GSSG) for 1 h at room temperature. Following incubation, unreacted dhG or GSSG was removed by dialysis against PBS. NISCH proteins were resolved by SDS-PAGE and subsequently transferred to PVDF membranes for immunoblotting. Protein glutathione modification or glutathionylation was assessed by probing with anti-glutathione antibody (Virogen, 101-A-250; 1:1000 dilution).

### Fusogenic liposome preparation

A stock solution of 1,2-dioleoyl-sn-glycero-3-phosphoethanolamine (DOPE), N,N,N-trimethyl-2,3-bis(oleoyloxy)propan-1-aminium methylsulfate (DOTAP), and 1,1’-dioctadecyl-3,3,3′,3’-tetramethylindotricarbocyanine iodide (DiR′) was prepared in chloroform at a weight ratio of 1:1:0.1 (w/w/w). The solvent was evaporated under vacuum to obtain a thin lipid film. The dried lipids (50 μg) were hydrated with 200 μL PBS (pH 7.4) containing 10 μg purified enNISCH protein, followed by vigorous mixing. The lipid suspension was sonicated on ice for 5 min and subsequently extruded through 0.1 μm pore-size Whatman Nuclepore Track-Etch membrane using an AVANTI Polar Lipids extruder. Protein-loaded liposomes were purified by gel filtration using NAP-5 columns (Cytiva).

### Protein delivery by fusogenic liposome

MDA-MB-231 cells were cultured in 10 cm dishes to approximately 70% confluency and washed once with PBS. Cells were maintained in DMEM for 30 min at 37°C prior to treatment. Fusogenic liposomes, encapsulating none or NISCH protein, were prepared in PBS (100 µL) and diluted with an equal volume of DMEM (100 µL). The mixtures were incubated for 1 h at room temperature before being added to the cells, followed by 3 h incubation at 37 °C. Cells were then washed four times with PBS (1 mL each). For validation of protein delivery, cells were lysed and analyzed by western blotting using anti-HIS antibody (Cell Signaling Cat #2365). Migration and invasion assays were performed as described above.

### Colony formation assay

MDA-MB-231 cells (500 cells/well) were seeded into 12-well plates. After 24 h, cells were serum-starved for 3 h, followed by treatment with varying concentrations of fusogenic liposomes containing enNISCH construct (0–40 µg/mL) prepared in serum-free DMEM (i.e., 25 mM glucose). The control cells were treated with a fusogenic liposome without encapsulating enNISCH. Treatment was applied for 3 h and repeated every 72 h. After 12 days, colonies were fixed with 4% paraformaldehyde, stained with 0.2% crystal violet, and washed extensively with PBS to remove excess dye. Colonies containing more than >50 cells were counted manually, and results were quantified and plotted.^79^

### Statistical analysis

All data are presented as means ± SD and were statistically analyzed using one-way ANOVA followed by Tukey’s *post-hoc* test, or two-way ANOVA followed by Tukey’s *post-hoc* test. The value p < 0.03 is statistically significant.

## Data availability

All data supporting the findings of this study are available from the corresponding author upon request.

## Supplementary information

This article contains supporting information.

## Acknowledgements

The research was supported by the National Institutes of Health (NIH) (R01 GM143214) (Y.H.A), a National Health and Medical Research Council (NHMRC) Investigator Grant (APP2016410) (B.M.C), and a research fund from Drexel University (Y.H.A). We would like to thank all the Ahn group members for assisting in the experiments.

## Author Contributions

Conceptualization, M.C.S., Y.H.A.; Formal Analysis, M.C.S., D.S.K.K., D.O., R.P., F.M.R., D.E., Y.H.A.; investigation, M.C.S., D.S.K.K., D.O., R.P., F.M.R.; supervision, B.C., Y.H.A.; visualization, M.C.S., D.S.K.K., D.O., R.P., F.M.R., Y.H.A.; writing – original draft, M.C.S., Y.H.A.; writing – review & editing, M.C.S., D.S.K.K., D.O., R.P., F.M.R., D.E., B.C., Y.H.A.; funding acquisition, Y.H.A.

