## Supplementary Information for "Redox Regulation of Cell Migration via Nischarin S-glutathionylation"

Shivamadhu *et al*.

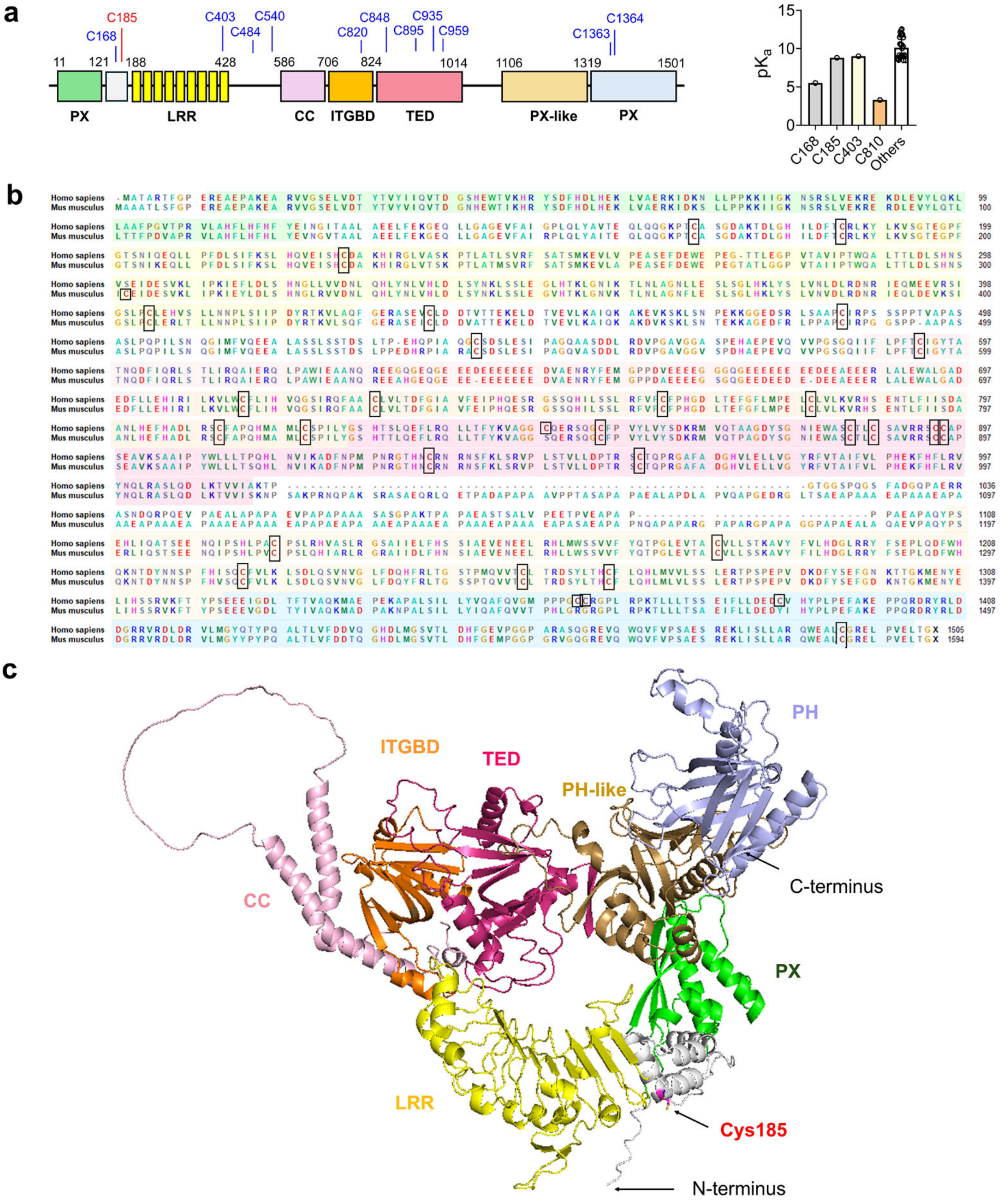


**Supplementary Figure 1. Analysis of NISCH structure, sequence, and Cys residues.** (**a**) NISCH domains and Cys residues. Among 32 cysteines, cysteines more exposed to the surface are indicated (left). Cysteine p$K_a$ analysis is shown (right). (**b**) NISCH sequence alignment between human and mouse. Cysteines are highlighted by boxes. (**c**) NISCH structure predicted by the AlphaFold. The structure was downloaded from UniProt. Domains are indicated by colors used in (**a**).

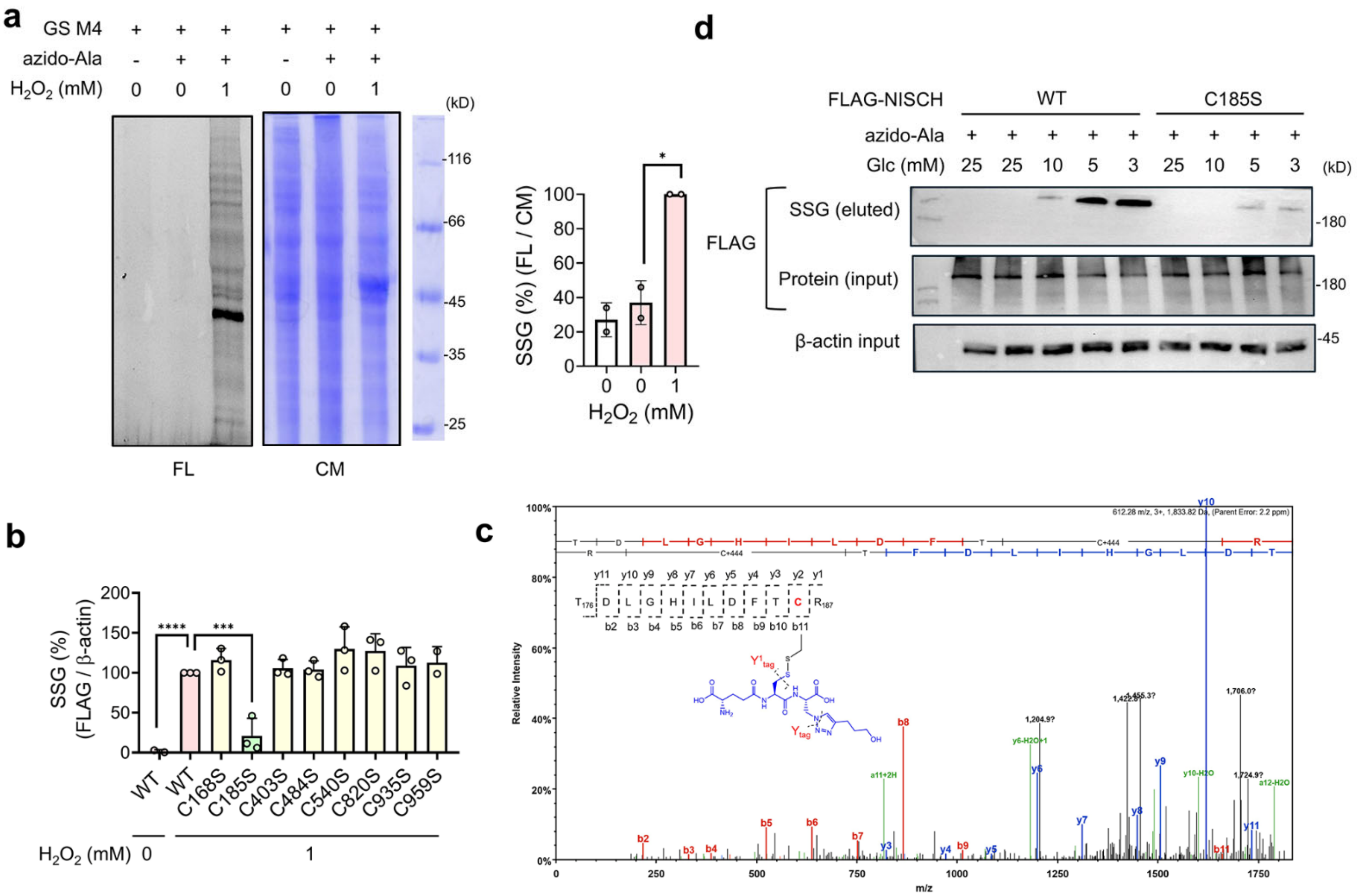


**Supplementary Figure 2. Glutathionylation analysis of NISCH in cells.** A clickable glutathione approach was used to monitor glutathionylation (SSG). Azido-Ala (0.6 mM) was incubated in HEK293/GS M4 cells or MDA-MB-231 cells expressing GS M4 for 20 h. After inducing glutathionylation, the lysates were collected and subjected to a click reaction with rhodamine-alkyne or biotin-alkyne. Glutathionylation was analyzed by fluorescence (FL) and Commassie stain (CM), or western blots before (input) and after (eluted) streptavidin-agarose pull-down. (**a**) Global glutathionylation in HEK293/GS M4 cells upon incubating with hydrogen peroxide (1 mM) for 15 min (n=2). (**b**) Glutathionylation of NISCH mutants. Quantification analysis of data in Figure 1F (n=3). (**c**) Tandem mass analysis of NISCH glutathionylation. The data was retrieved from our previous proteomic data,[S1] detecting mouse NISCH C186 glutathionylation (from mouse HL-1 cell line). Note that human NISCH C185 is equivalent to mouse NISCH C186: please see Figure S1B. (**d**) Glutathionylation of NISCH WT and C185S mutant. FLAG-NISCH WT or C185S was transiently transfected into MDA-MB2-231 cells. Glutathionylation was induced in different concentrations of glucose (Glc) (3-25 mM). Data represent the mean ± SD. The statistical difference was analyzed by one-way ANOVA and Tukey's post-hoc test (**a**-**b**), where *p < 0.03, **p < 0.002, ***p < 0.0002, ****p < 0.0001.

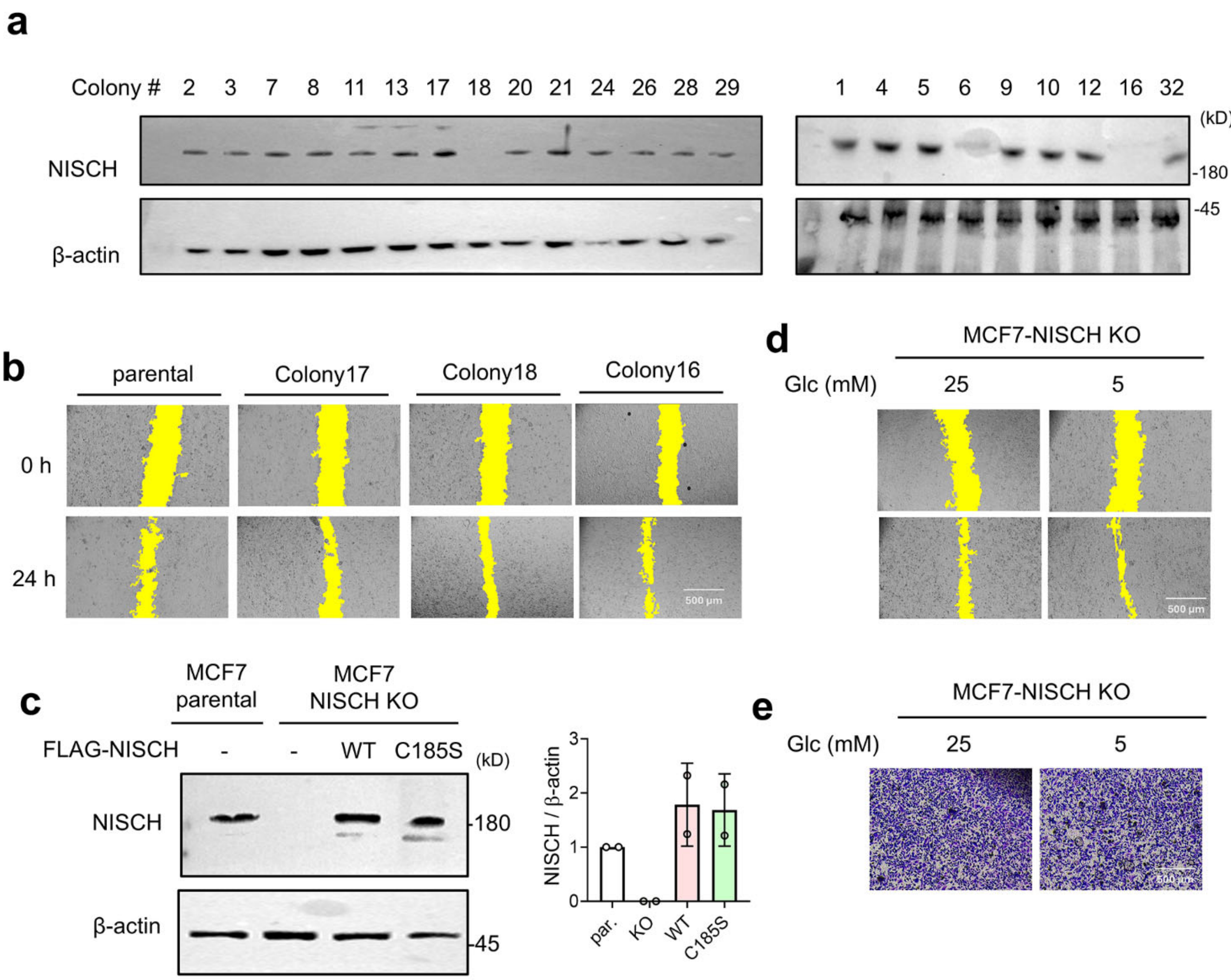


**Supplementary Figure 3. Construction and evaluation of MCF7 NISCH knockout cell line.** (**a**) Analysis of colonies of MCF7 cells after NISCH knockout (KO). After transfection of CRISPR/Ca9 plasmids, cells were split into individual clones and grown into colonies, which were analyzed by western blot. (**b**) In vitro scratch migration analysis of selected colonies (n=3). A scale bar = 500 μm. Quantification analysis is shown in Figure 2A. (**c**) NISCH re-expression into MCF7 NISCH KO cells (colony 18). After transfection of FLAG-NISCH WT or C185S, cells were analyzed by western blot (n=2). (**d**) In vitro scratch migration analysis of MCF7-NISCH KO cells (colony 18) (n=6). A scale bar = 500 μm. The quantification analysis is shown in Figure 2b. (**e**) Transwell migration analysis of MCF7-NISCH KO cells (n=2). A scale bar = 500 μm. The quantification analysis is shown in Figure 2d. In **b**-**d**, yellow colors indicate the area without cells. Data represent the mean ± SD. The statistical difference was analyzed by one-way ANOVA and Tukey's post-hoc test (**c**), where *p < 0.03, **p < 0.002, ***p < 0.0002, ****p < 0.0001.

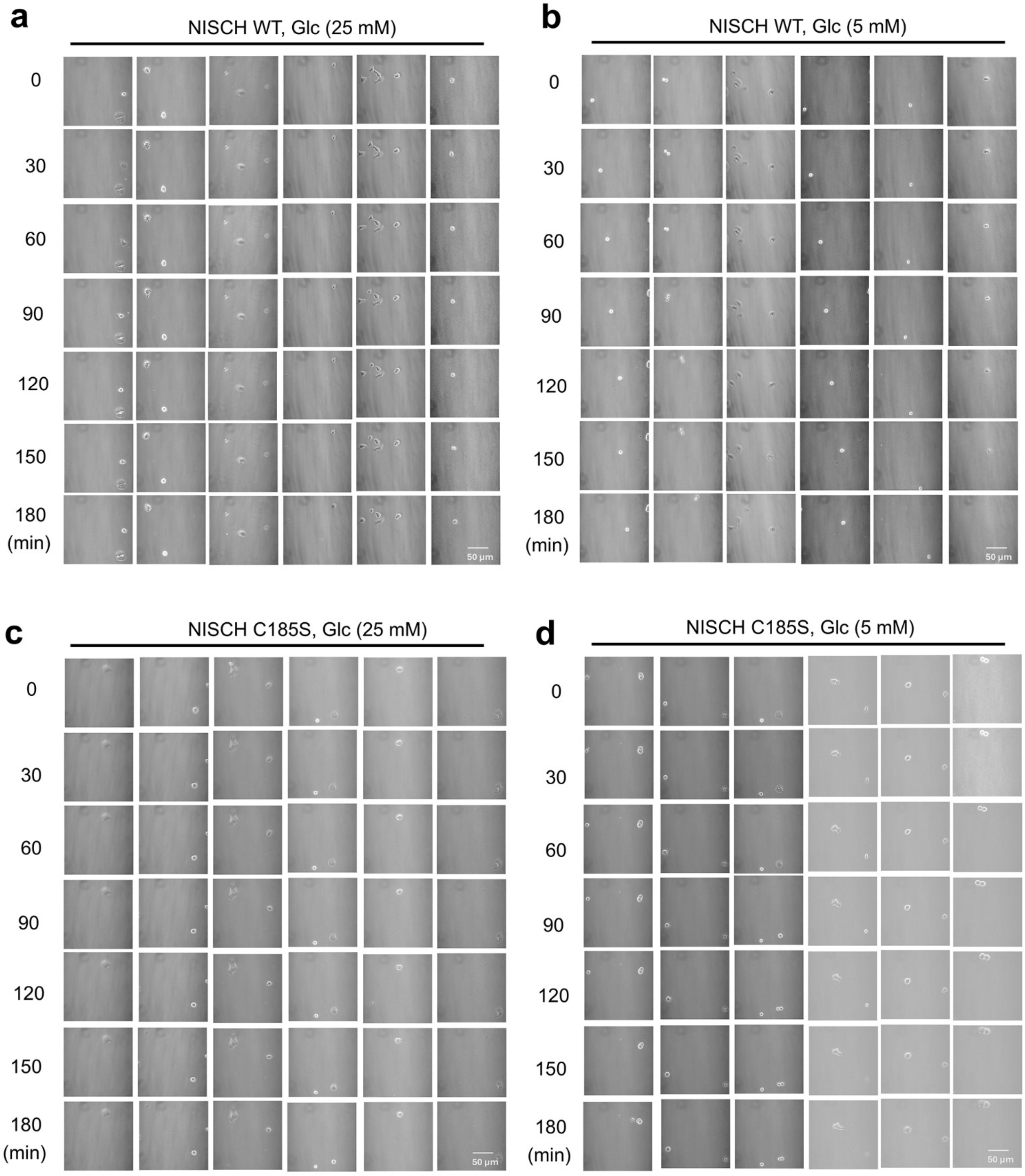


**Supplementary Figure 4. Migration analysis of single cells expressing NISCH WT or C185S.** MCF7-NISCH KO cells were transfected with FLAG-NISCH WT or C185S and selected in the presence of G418. Cells were plated at low confluency, incubated in high or low glucose conditions, and monitored under a confocal microscope for 3 h. Images were taken every 10 min (n = 10 cells in 6 images). A scale bar = 50 μm. The representative images and quantification analysis are shown in Figure 2c.

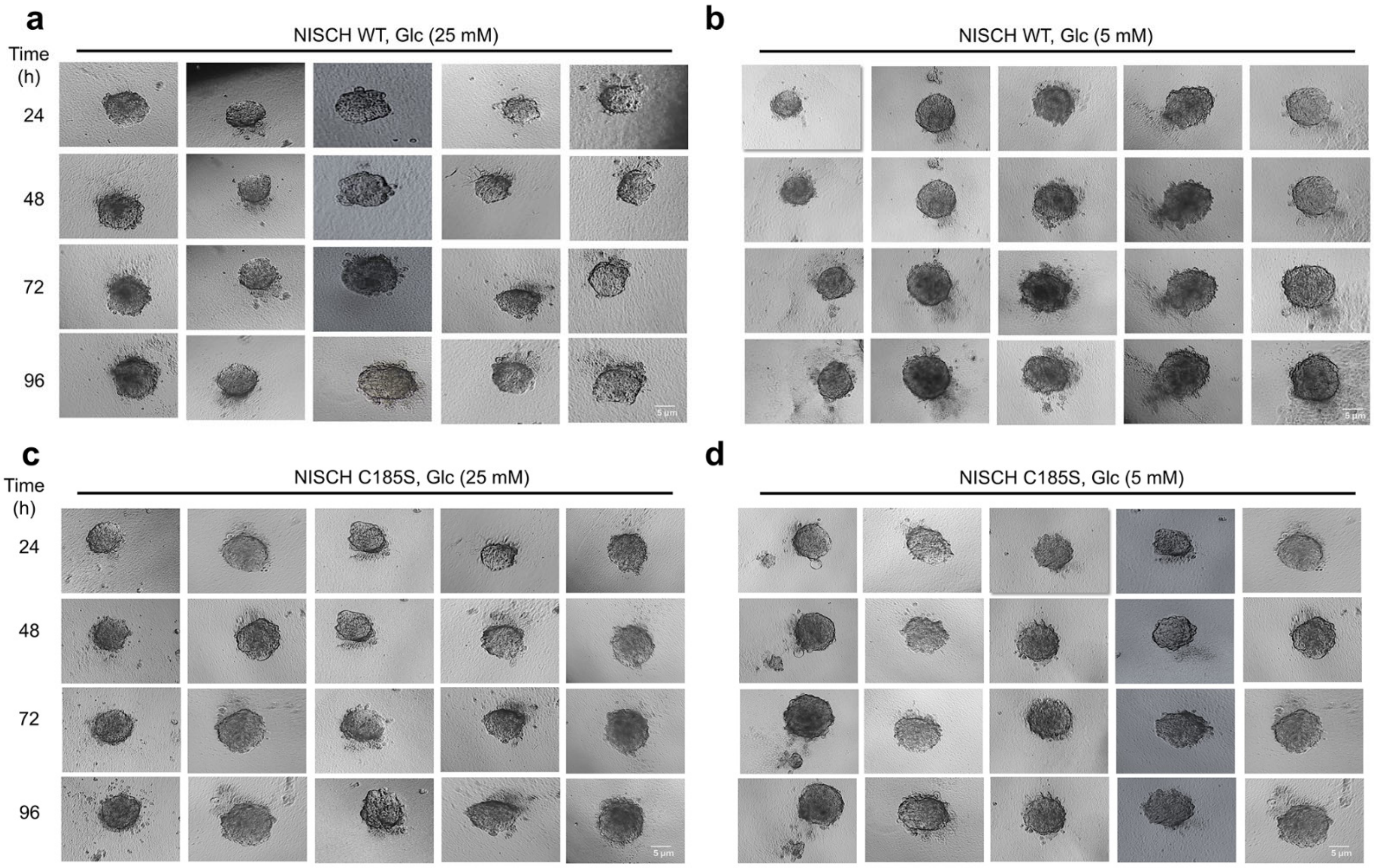


**Supplementary Figure 5. Spheroid invasion assay of MCF7 cells expressing NISCH WT or C185S.** MCF7-NISCH KO cells were transfected with FLAG-NISCH WT or C185S and selected in the presence of G418 (100 µg/mL). After forming spheroids, individual spheroids were plated in 96-well plates in Matrigel:collagen I (1:1) and incubated in high or low glucose conditions, and monitored for invading cells into the matrix over 96 h. Images show replicate data in Figure 2e, and the quantification analysis is shown in Figure 2e.

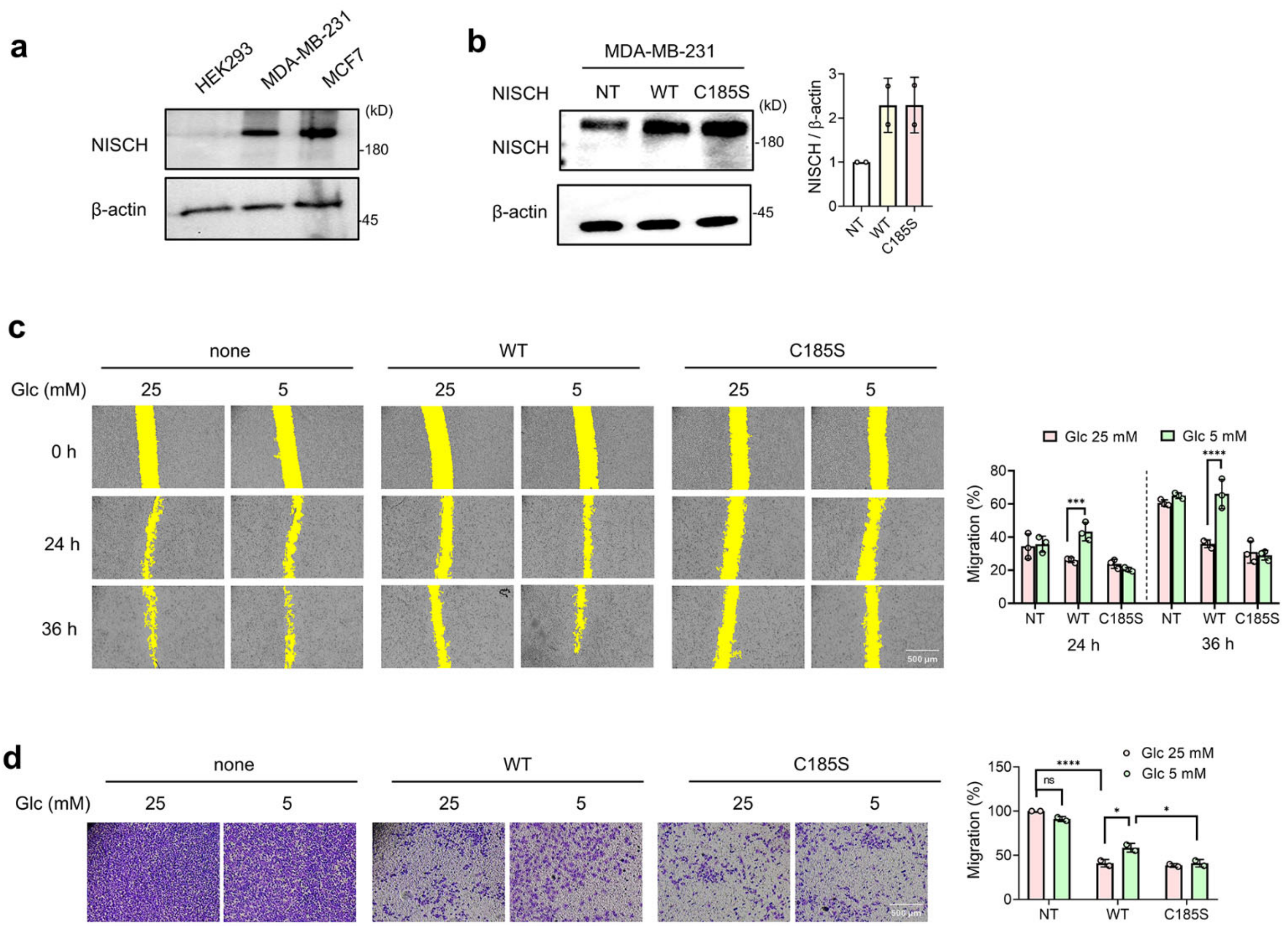


**Supplementary Figure 6. Migration and invasion analysis of MDA-MB-231 cells expressing NISCH WT or C186S.** MDA-MB-231 cells were transfected with NISCH WT or C185S. (**a**) Endogenous expression level of NISCH in cell lines. (**b**) NISCH expression levels after ectopic expression of FLAG-NISCH in MDA-MB-231 cells. NT= no transfection. (**c**) In vitro scratch migration analysis of MDA-MB-231 cells expressing NISCH WT or C185S. After transfection, cells were incubated in high (25 mM) or low (5 mM) glucose conditions (n=3). A scale bar = 500 µm. Yellow colors indicate the area without cells. (**d**) Transwell invasion analysis. After transfection of NISCH constructs, cells were incubated in high (25 mM) or low (5 mM) glucose conditions for 24 h (n=2). A scale bar = 500 µm. Data represent the mean ± SD. The statistical difference was analyzed by one-way ANOVA (**b**) or two-way ANOVA (**c**-**d**) with Tukey's post-hoc test, where *p < 0.03, **p < 0.002, ***p < 0.0002, ****p < 0.0001.

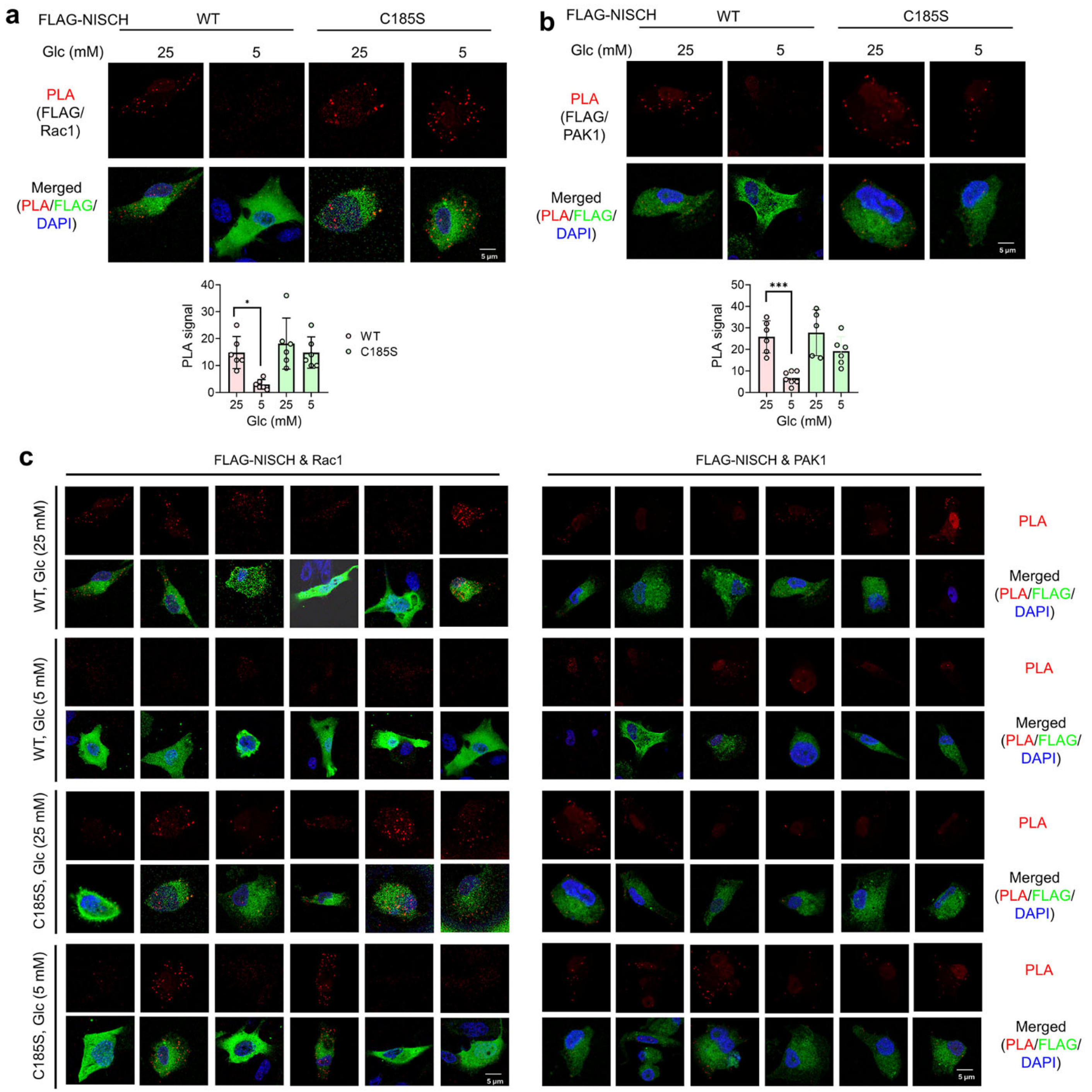


**Supplementary Figure 7. Proximity ligation analysis of NISCH and its binding partners.** MDA-MB-231 cells expressing FLAG-NISCH WT or C185S were incubated in high (25 mM) or low (5 mM) glucose conditions for 24 h. After fixation, cells were analyzed by proximity ligation assay (PLA) (red), immunostaining using FLAG-antibody (NISCH, green), and DAPI (blue)**.** (**a**) PLA using antibodies toward FLAG-NISCH and Rac1 (n= 6 images). A scale bar = 5 µm. (**b**) PLA using antibodies toward FLAG-NISCH and PAK1 (n=6 images). A scale bar = 5 µm. (**c**) Replicate images of data in **a**. and **b**. Data represent the mean ± SD. The statistical difference was analyzed by one-way ANOVA and Tukey's post-hoc test (**a**, **b**), where *p < 0.03, **p < 0.002, ***p < 0.0002, ****p < 0.0001.

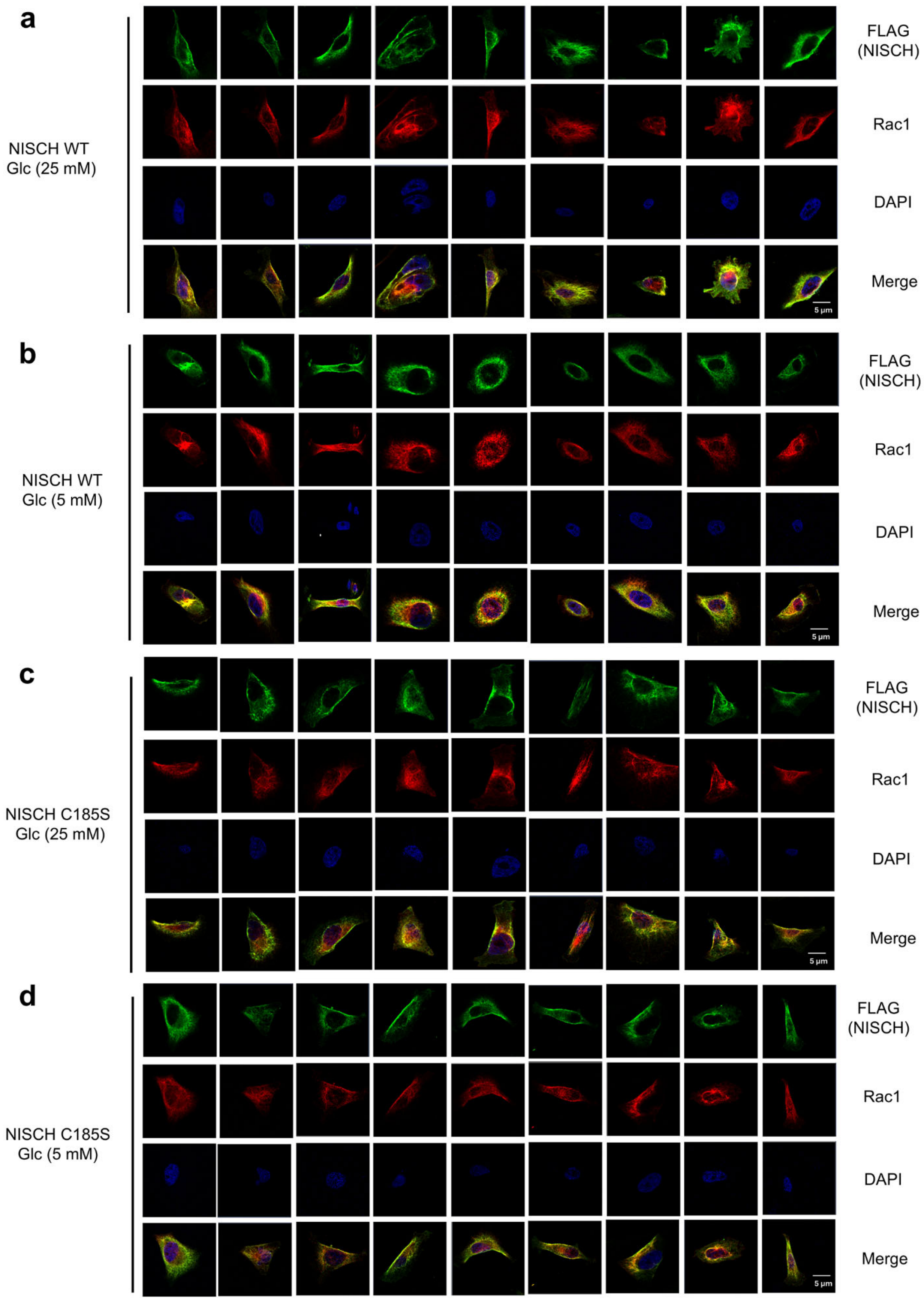


**Supplementary Figure 8. Co-localization analysis of NISCH and Rac1.** MDA-MB-231 cells expressing FLAG-NISCH WT or C185S were incubated in high (25 mM) or low (5 mM) glucose conditions for 24 h. Replicate images are shown and include the data in Figure 3d. The representative images and quantification analysis are shown in Figure 3d. A scale bar = 5 µm.

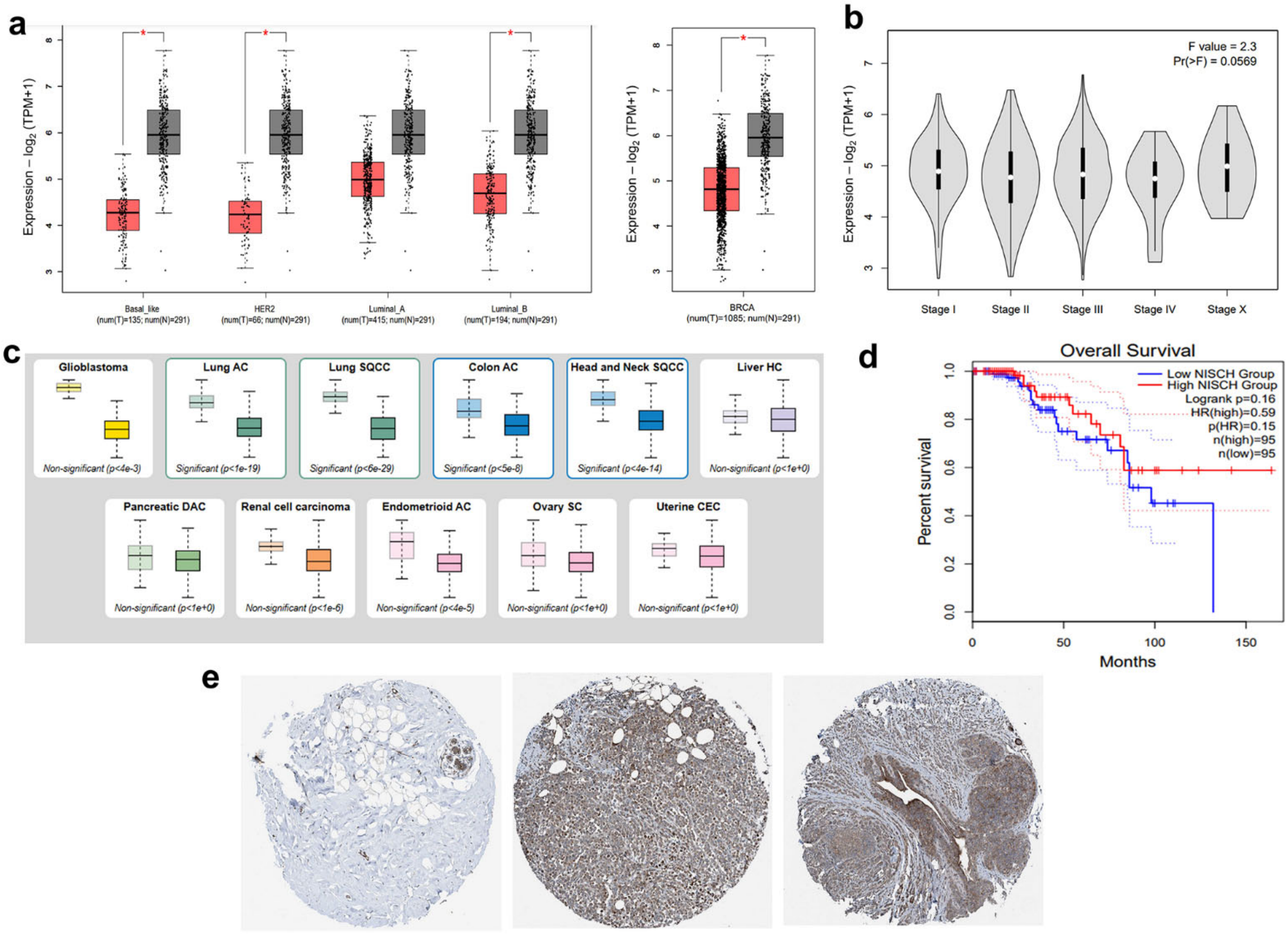


Breast
HPA023189
Female, age 23
Breast (T-04000)
Normal tissue. NOS (M-00100)
Patient ID: 2883

Breast Cancer
HPA023189
Female, age 61
Breast (T-04000)
Duct carcinoma
(M-085003)
Patient ID: 1910

Breast Cancer
HPA023189
Female, age 50
Breast (T-04000)
Lobular carcinoma
(M-085203)
Patient ID: 2407

**Supplementary Figure 9. Transcriptomic and protein expression analysis of NISCH in cancer.** Transcriptomic and proteomic datasets were analyzed to evaluate the role of NISCH in cancer (Tumor: 1085, Normal: 291). (**a-b**) NISCH expression in different types (**a**) and stages (**b**) of breast cancer. RNA-seq expression data were obtained from The Cancer Genome Atlas (TCGA) and the Genotype-Tissue Expression (GTEx) project through the GEPIA2 platform. Differential expression of NISCH between tumor and normal tissues was assessed using GEPIA2 across multiple cancer types. Subgroup analyses were performed for breast cancer molecular subtypes (Basal, HER2+, Luminal A, Luminal B) and tumor stages. Expression is shown with the unit of transcripts per million (TPM). (**c**) NISCH expression levels in different cancers. Protein-level expression was retrieved from the Human Protein Atlas (HPA). (**d**) The correlation of survival versus NISCH expression. Kaplan–Meier survival curves were generated using GEPIA2 to examine the relationship between NISCH expression and overall survival (OS) in breast cancer patients. Median expression was used as the cutoff to define high and low groups. Hazard ratios (HR) and log-rank p-values were calculated. (**e**) NISCH immunohistochemistry (IHC) in breast cancer tissues. Representative IHC images for NISCH in normal and tumor tissues were obtained from the Human Protein Atlas (HPA). Expression levels and localization were compared qualitatively.

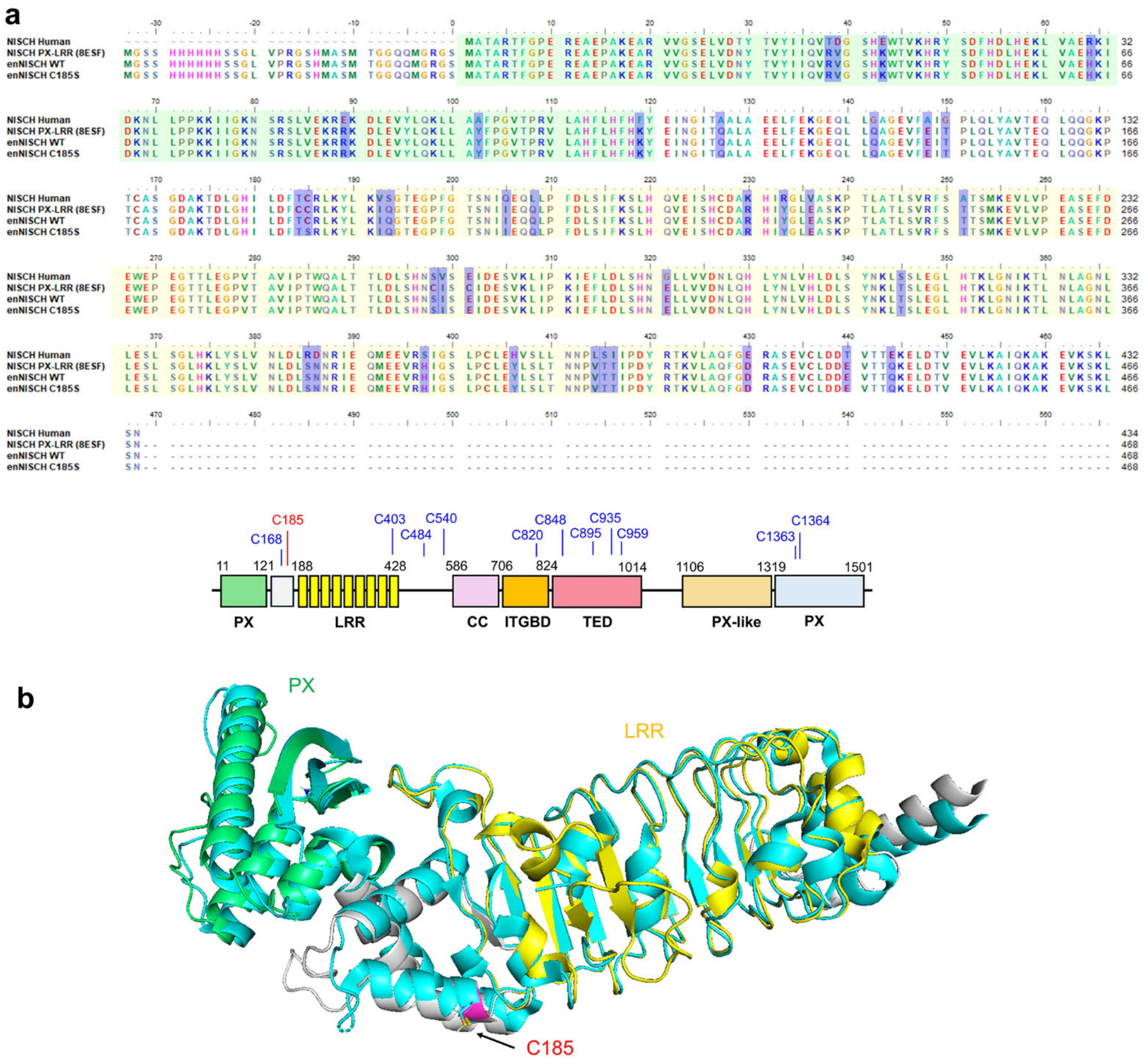


**Supplementary Figure 10. The engineered NISCH construct.** (**a**) The sequence comparison of NISCH constructs: 1) NISCH WT, 2) The engineered NISCH N-terminal domain encompassing PX-LRR (NISCH PX-LRR). This is the construct for the X-ray crystal structure reported in the Protein Data Bank (PDB: 8ESF), 3) The engineered NISCH by introducing additional mutations (C184T, C298S, and C300E) into NISCH PX-LRR (enNISCH WT). 4) The engineered NISCH by introducing additional mutations (C184T, C298S, C300E and C185S) into NISCH PX-LRR (enNISCH C185S). (**b**) The structure overlap between NISCH WT (AlphaFold) and NISCH PX-LRR (PDB: 8ESF). The Alpha-Fold structure is shown in colors representing domains (i.e., PX: green, A region between PX and LRR: white, LRR: yellow). The NISCH-PX-LRR is shown in cyan.

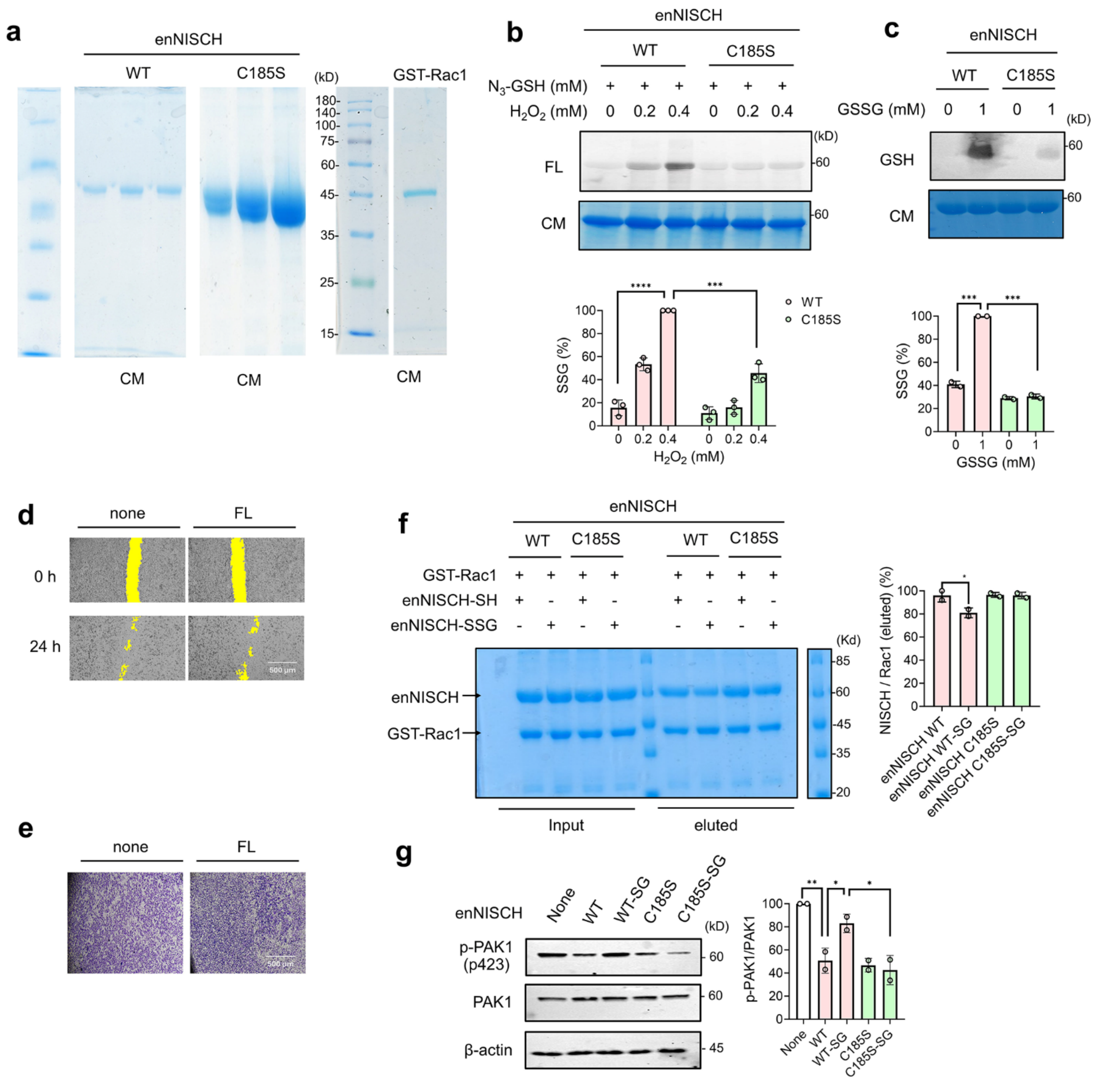

**Supplementary Figure 11. Analysis of enNISCH in vitro and in cells.** (**a**) The purity of recombinant enNISCH constructs and GST-Rac1. (**b-c**) In vitro glutathionylation analysis of enNISCH WT and C185S. Purified enNISCH WT or C185S was incubated with azido-glutathione and hydrogen peroxide for 15 min, subjected to a click reaction with rhodamine-alkyne, and analyzed by in-gel fluorescence (FL) or Coomassie stain (CM) (n=3) (**b**). Purified enNISCH WT and C185S were incubated with oxidized glutathione (GSSG) for 1 h and analyzed by western blot using glutathione antibody (GSH) or Coomassie stain (CM) (n=2) (**c**). (**d-e**). In vitro scratch migration (**d**) and transwell invasion (**e**) assays of MDA-MB-231 cells upon incubating with fusogenic liposomes for 2 h. Images are the representative data shown in Figures 5d and 5e. A scale bar = 500 μm. Quantification analyses are shown in Figures 5d and 5e. (**f**) In vitro analysis of enNISCH binding interaction with GST-Rac1. GST-Rac1 was immobilized to glutathione-agarose, then incubated with enNISCH pre-incubated with none (enNISCH-SH) or GSSG (enNISCH-SSG). The eluted samples were analyzed in SDS-PAGE with Commassie stains (CM) (n=2). (**g**) Analysis of enNISCH for its effects on PAK1 in cells. MDA-MB-231 cells were incubated with fusogenic liposome containing enNISCH constructs (enNISCH WT, enNISCH WT-SG, enNISCH C185S, and enNISCH C185S-SG: glutathione modification by dhG) for 2 h, and the lysates were analyzed by PAK1 phosphorylation by western blot (n=2). Data represent the mean ± SD. The statistical difference was analyzed by one-way ANOVA and Tukey's post-hoc test (**b-c, f-g**), where $^*p < 0.03$, $^{**}p < 0.002$, $^{***}p < 0.0002$, $^{****}p < 0.0001$.

| | | |
|---|---|---|
| FLAG-NISCH site-directed mutagenesis | | |
| C185S | Forward | CCT GGA CTT CAC CTC TCG CCT TAA GTA CC |
| | Reverse | GGT ACT TAA GGC GAG AGG TGA AGT CCA GG |
| | Forward | TCC AGC AGG GTA AAC CCA CCT CCG CGT CAG GTG ACG CAA AG |
| C168S | Reverse | CTT TGC GTC ACC TGA CGC GGA GGT GGG TTT ACC CTG CTG GA |
| C403S | Forward | TCT ATA GGA AGC CTT CCA TCT TTG GAA CGA CTG ACC C |
| | Reverse | GGG TCA GTC GTT CCA AAG ATG GAA GGC TTC CTA TAG A |
| C484S | Forward | TCC GGC CCC CTG CAT TCG GCC CGG AGG TTC C |
| | Reverse | GGA ACC TCC GGG CCG AAT GGA GGG GG CC GGA |
| C540S | Forward | ATC GCC AGG GCT TCC AGT GAC TCT CTC GAG AGT ATT CC |
| | Reverse | GGA ATA CTC TCG AGA GAG TCA CTG GAA GCC CTG GCG AT |
| C820S | Forward | AAC ACA TGG CCA TGC TGT CCA GCC CTA TCC TCT ACG GAT C |
| | Reverse | GAT CCG TAG AGG ATA GGG CTG GAC AGC ATG GCC ATG TGT T |
| C935S | Forward | AAT AGA GGG ACA CAT AAC TCT CGG AAC CGG AAC TCT TTC AAG C |
| | Reverse | GCT TGA AAG AGT TCC GGT TCC GAG AGT TAT GTG TCC CTC TAT T |
| C959S | Forward | TCG ACC CTA CTA GGA GCT CCA CCC AGC CCA G |
| | Reverse | CTG GGC TGG GTG GAG CTC CTA GTA GGG TCG A |
| enNISCH site-directed mutagenesis | | |
| C184T | Forward | CAT ATT CTG GAC TTC ACC TGT CGC CTG AAA TAT CTG AAA ATTC |
| | Reverse | GAA TTT TCA GAT ATT TCA GGC GAC AGG TGA AGT CCA GAA TAT G |
| C185S | Forward | CAT ATT CTG GAC TTC AGC TGT CGC CTG AAA TAT CTG AAA ATT C |
| | Reverse | GAA TTT TCA GAT ATT TCA GGC GAC AGC TGA AGT CCA GAA TAT G |
| C298S | Forward | CTG AGC CAT AAT TCC ATT AGC TGC ATT GAT G |
| | Reverse | CAT CAA TGC AGC TAA TGG AAT TAT GGC TCA G |
| C300E | Forward | CCA TAA TTG CAT TAG CTC CAT TGA TGA AAG CGT G |
| | Reverse | CAC GCT TTC ATC AAT GGA GCT AAT GCA ATT ATG G |
| CRISPR-KO-primers | | |
| gRNA1 | Sense | CACCG ATC AAT ATT CAA GTC CCT GC |
| | Antisense | AAA GCA GGG ACT TGA ATA TTG AT |
| gRNA2 | Sense | CACCG AAC ATC AAG ACC TTA AAC C |
| | Antisense | AAA GGT TTA AGG TCT TGA TGT TC |
| gRNA3 | Sense | CACCG GGC CAA GGA AGC GCG CGT CG |
| | Antisense | AAAC CGA CGC GCG CTT CCT TGG CC |

**Supplementary Table 1. A summary of cloning primers.**

**Additional Methods**

**Proximity Ligation Assay.** To examine protein–protein interactions in situ, Duolink® Proximity Ligation Assay (PLA; Sigma-Aldrich) was performed according to the manufacturer's instructions with minor modifications. NISCH WT and C185S transfected MDA-MB-231 cells were seeded in a 35 mm glass-bottom dish coated with fibronectin (0.1%). After incubating 5 mM or 25 mM glucose conditions, cells were fixed with 4% paraformaldehyde (500 µL) at room temperature for 10 min, then permeabilized with 0.2% Triton X-100 for 10 min. After blocking with Duolink blocking buffer for 1 h at 37 °C, cells were incubated overnight at 4 °C with primary antibody pairs raised in different species against PAK1 (1:200; Cell Signaling, Cat #2608S) and Rac1/cdc42 (1:200; Cell Signaling, Cat #4651S), and FLAG (NISCH, 1:200; Milipore-Sigma, Cat# F3165). Negative controls included samples incubated with only one primary antibody or with species-matched IgG controls. Following primary antibody incubation, samples were washed and incubated with species-specific PLA probes (anti-rabbit PLUS and anti-mouse MINUS) for 90 mins at 37 °C. The ligation reaction was carried out by diluting Duolink ligation buffer in distilled water (1:5 dilution), followed by immediately adding ligase (1:40 dilution). The ligation mixture was incubated at 37°C in a humidified incubator for 30 min. Following ligation, cells were washed twice with 1× Duolink wash buffer A for 5 min each. The amplification polymerase reaction was then performed by diluting the amplification stock in distilled water (1:5 dilution) and adding polymerase (1:80 dilution) at the time of incubation. Cells were incubated with the amplification mixture for 100 min at 37°C, after which they were washed twice with 1× Duolink wash buffer B for 15 min each. To detect FLAG expression, cells were incubated with anti-mouse Alexa Fluor 488 (1:1000; Invitrogen, Cat# A-21235) for 1 h at room temperature. Following antibody incubation, cells were washed three times with 1× Duolink wash buffer B and once with 0.01× Duolink wash buffer B before being mounted with Prolong™ Gold Antifade mounting DAPI (Invitrogen, Cat #P36931) for 10 min at room temperature. Fluorescent images were acquired using a Zeiss LSM700 inverted confocal microscope, and data were analyzed using ImageJ software. Filter settings were adjusted for DAPI to detect nuclear staining (excitation wavelength 350 nm, emission wavelength 470 nm), Texas Red for PLA puncta (excitation wavelength 594 nm, emission wavelength 624 nm), and Alexa Fluor 488 for NISCH-FLAG detection (excitation wavelength 490 nm, emission wavelength 525 nm).

Images were first split into individual channels (Image > Color > Split Channels). The red channel, corresponding to the PLA signal, was used for quantification, while the merged images (red channel + DAPI) were generated for visualization. Thresholds were manually adjusted (Image > Adjust > Threshold) with "Dark background" selected, to isolate puncta from background noise. Puncta were quantified using the "Analyze Particles" function (Analyze > Analyze Particles), with particle size set between 0.2–5 $\mu m^2$ to exclude noise and artifacts. Parameters selected included *Display Results*, *Summarize*, and *Exclude on Edges*, thereby generating particle counts, area, and spatial localization. Data were interpreted as the average number of PLA puncta per cell, the percentage of PLA-positive cells, and, in some cases, total fluorescence intensity.

**In vitro binding analysis of enNISCH and GST-Rac1**. 20 µg of purified His-tagged NISCH N-terminal domain in the presence or absence of 1 mM GSSG for 1 h at room temperature under constant rotation, and excess glutathione was removed using dialysis for 3 h. On the other hand, purified GST-Rac1 (10 µg) was incubated in binding buffer (20 mM Tris-HCl, pH 7.5, 50 mM NaCl, 5 mM EDTA) supplemented with 100 µM GTPγS (Milipore Sigma # 20-176) at room temperature for 30 min. The reaction was terminated by the addition of $MgCl_2$ (final concentration 10 mM) and further incubated with prewashed 20 µL glutathione-Sepharose beads (Cytiva #17513201) and incubated for 30 min at room temperature under gentle rotation. GST-Rac1-GTP complexes on beads were then incubated with 20 µg of purified His-tagged enNISCH-SH or enNISCH-SG under gentle rotation. After incubation, beads were washed extensively with wash buffer (20 mM Tris-HCl, pH 7.5, 50 mM NaCl, 5 mM EDTA, 0.1% Triton X-100) to remove unbound proteins. Bound proteins were eluted by boiling in SDS sample buffer (10% SDS) at 95°C for 5 min, resolved by SDS–PAGE, and images were documented.
